# Complementary Cytoskeletal Feedback Loops Control Signal Transduction Excitability and Cell Polarity

**DOI:** 10.1101/2024.02.13.580131

**Authors:** Jonathan Kuhn, Parijat Banerjee, Andrew Haye, Douglas N. Robinson, Pablo A. Iglesias, Peter N. Devreotes

## Abstract

To move through complex environments, cells must constantly integrate chemical and mechanical cues. Signaling networks, such as those comprising Ras and PI3K, transmit chemical cues to the cytoskeleton, but the cytoskeleton must also relay mechanical information back to those signaling systems. Using novel synthetic tools to acutely control specific elements of the cytoskeleton in *Dictyostelium* and neutrophils, we delineate feedback mechanisms that alter the signaling network and promote front- or back-states of the cell membrane and cortex. First, increasing branched actin assembly increases Ras/PI3K activation while reducing polymeric actin levels overall decreases activation. Second, reducing myosin II assembly immediately increases Ras/PI3K activation and sensitivity to chemotactic stimuli. Third, inhibiting branched actin alone increases cortical actin assembly and strongly blocks Ras/PI3K activation. This effect is mitigated by reducing filamentous actin levels and in cells lacking myosin II. Finally, increasing actin crosslinking with a controllable activator of cytoskeletal regulator RacE leads to a large decrease in Ras activation both globally and locally. Curiously, RacE activation can trigger cell spreading and protrusion with no detectable activation of branched actin nucleators. Taken together with legacy data that Ras/PI3K promotes branched actin assembly and myosin II disassembly, our results define front- and back-promoting positive feedback loops. We propose that these loops play a crucial role in establishing cell polarity and mediating signal integration by controlling the excitable state of the signal transduction networks in respective regions of the membrane and cortex. This interplay enables cells to navigate intricate topologies like tissues containing other cells, the extracellular matrix, and fluids.

## Introduction

Cell migration is a fundamental process that plays critical roles in development, immune system function, and disease progression, including metastasis. While there is a broad diversity of cell migration strategies, efficient motion usually depends on the spontaneous formation of membrane protrusions and the establishment of front-back polarity. Both protrusion formation and polarity rely on coordination between actin cytoskeletal dynamics and signaling networks. The precise mechanisms by which signaling controls cytoskeletal activity are still being unraveled, and even less is known about the potential feedback from changes in cytoskeletal organization and mechanics on these same signaling networks^1,2^.

Regulation of the actin cytoskeleton is intricately linked to signaling molecules, such as Ras GTPases and phosphoinositides. The activation of Ras GTPase, associated signaling molecules, and the lowering of anionic lipids correlates strongly with the formation of cell protrusions^3–8^. In amoeboid migrating cells, protrusion formation involves the activation of branched actin nucleator Arp2/3 and the local disassembly of myosin II (hereafter referred to as ‘myosin’) filaments and actin-actin crosslinkers^9–14^. Previous studies have suggested positive feedback exists between Arp2/3-mediated actin polymerization and PI3K activity at the front^8,11,15–18^. Synergistically, there is evidence that negative feedback from actomyosin inhibits signaling. For example, cells lacking myosin have elevated Ras activation, and retracting pseudopods are insensitive to chemotactic stimulation^19–21^. Together these findings suggest that branched actin and actomyosin networks may differentially feedback on signaling networks. However, because the network governing cell migration is complex and adaptable, local and acute perturbations are critical to dissect the cause and nature of cytoskeletal feedback.

Both signal transduction networks and actin cytoskeletal networks exhibit properties of excitability, including an activation threshold and propagating waves^22–29^. Waves of Ras-GTP, phosphatidylinositol 3,4,5-trisphosphate (PIP3), actin polymerization, and Arp 2/3 traverse the basal cell membrane and define the “front” state of the cortex. Complementary zones (“shadow waves”) of PI(4,5)P2, cortexillin, and myosin II depletion define the “back” state of the cortex. While waves of signaling and cytoskeletal activity are coupled, they are morphologically distinct and independent^24^. In the absence of the cytoskeleton, signaling waves still propagate but cannot create protrusions, and in the absence of signaling activity, cytoskeletal activity exhibits excitable properties but does not propagate in broad waves that move cells^22,27^. These networks have been referred to as the Signal Transduction Excitable Network (STEN) and the Cytoskeletal Excitable Network (CEN), respectively. Previous studies suggest that modifying the threshold of STEN tunes the size and speed of these waves and ultimately the nature of actin-based protrusions^3,24^. Changes in wave behavior are therefore a surrogate for alterations in the STEN and CEN and can predict effects on cell behavior.

The properties of the signaling and cytoskeletal networks can be captured in computational models ^3,24,30–33^. Excitable systems require both fast positive and delayed negative feedback^34^. In the models, we have previously reported mutual inhibition between front and back activities, which leads to a positive feedback loop, and molecules at the cell front produced their own inhibitor on longer timescales^3,22^. These models predicted the behavior of coupled STEN-CEN waves and the response of cells to chemotactic stimulus. However, these models oversimplified the behavior of the cytoskeleton in this process. CEN activity represented the activation of Arp2/3 by Rac at the cell front and did not consider regulation to and feedback from the morphologically distinct actomyosin network at the cell back^24^. Furthermore, they have not been able to easily explain experimental observations about front-back polarity, which was shown to be dependent on cytoskeletal architecture^35^.

We sought to understand the impact of modulating the F-actin or actomyosin cytoskeletal structures on the STEN. First, we uncovered a positive feedback mechanism that promotes the front state. By increasing the pool of available actin monomers and promoting the formation of branched actin networks, STEN activation is enhanced. Second, it was shown previously that increasing Ras activity can lower myosin assembly^12,36^. Here, by designing precise chemically induced dimerization system (CID) and optogenetic tools to control myosin assembly in real time, we found that disassembly of the actomyosin cortex increases STEN activity. Together, these mutually negative effects constitute a positive loop that augments the back state.

Consistently, increasing the abundance of linear actin and crosslinked actin inhibits STEN activation. Moreover, inhibiting branched actin nucleation further enhances the overall effect. Our study here delineates the reciprocal relationship between cytoskeletal dynamics and signaling networks that takes place during cell migration. The cross-interaction of these molecular mechanisms allow cells to integrate signal transduction and mechanical cues and successfully navigate complex three-dimensional environments.

## Results

### Positive and negative feedback loops from the cytoskeleton control signaling networks

To investigate the effects of altering the cytoskeleton on signaling, we first examined the impact of increasing free monomeric actin in *Dictyostelium*, a classic model of amoeboid migration. Previously we found that mutating actobindins, which bind and sequester free G-actin, increases actin polymerization at the cell front^37,38^. To determine the effects of this on signaling, we measured Ras activation levels in *actobindin* triple knockout (*ABN ABC-*) cells to examine the effect of increased “front” actin network formation on STEN activity (**Fig. 1A-C, Movie S1**). Significantly, the *ABN ABC-* cells exhibited elevated levels of Ras activation, suggesting a positive regulatory role for front actin networks in STEN. To assess whether this increase is mediated by enhanced Arp2/3 activity, we measured the localization of WAVE complex subunit HSPC300 to the cell front in *ABN ABC-* and wild-type cells. As hypothesized, *ABN ABC-* cells showed greater recruitment of HSPC300 to the cortex compared to controls (**Fig. S1**). This result indicates the presence of a positive feedback loop from the front actin, which appears to form a branched Arp2/3-assembled network, to STEN.

**Figure 1:**
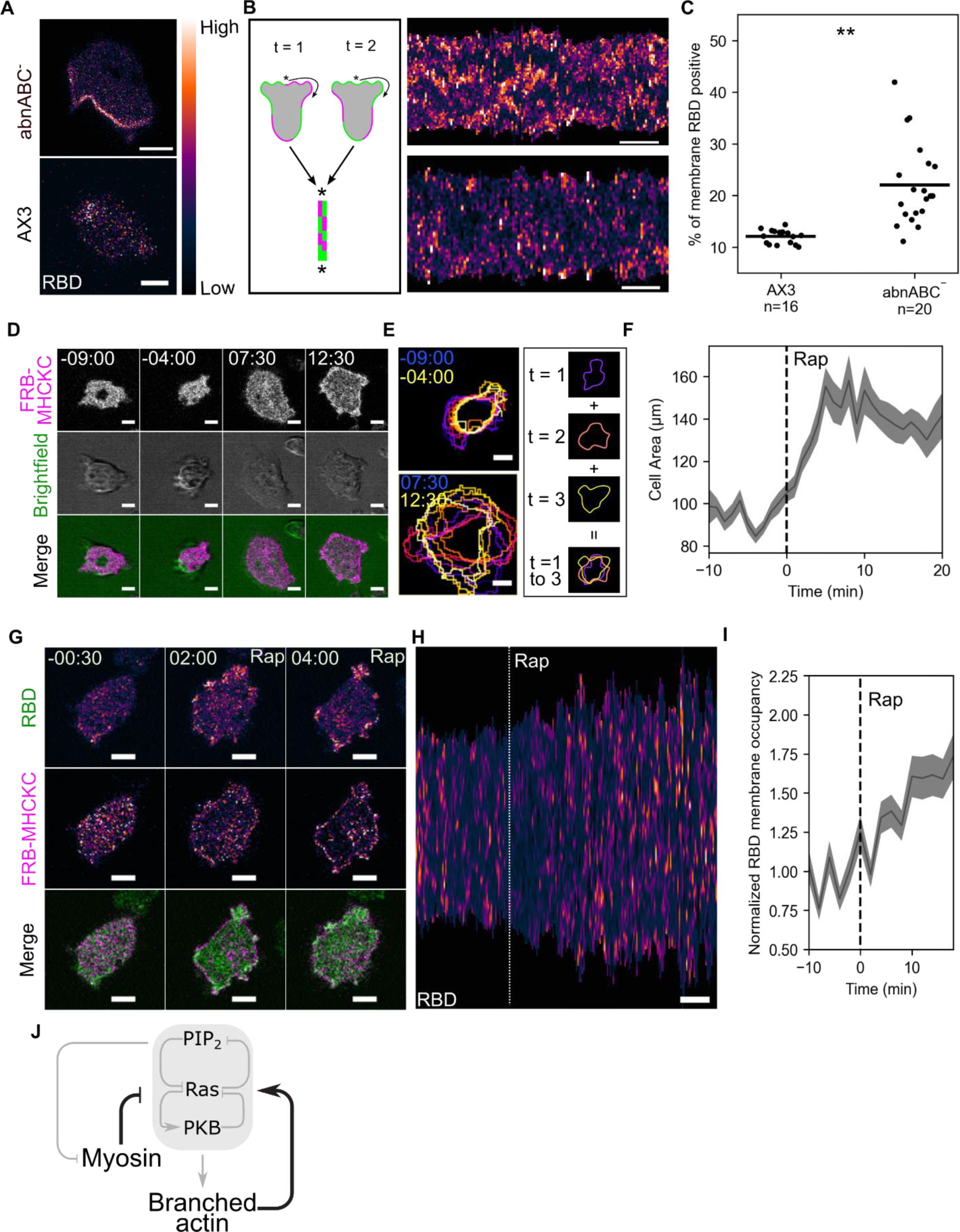
Cytoskeletal feedback loops at the cell front and back control Ras activity. **(A)** Scanning confocal imaging of Ras activation (RBD-EGFP) in wild-type (AX3) or *actobindin ABC* triple knockout (*abnABC^-^*) *Dictyostelium* cells. **(B)** Diagram (left) and experimental data (right) of RBD membrane kymographs corresponding to the movies in (A) (right). *Scale bars = 2 minutes*. **(C)** Individual (dots) and average (lines) percentages of the membrane periphery with RBD localization significantly above background in AX3 and *abnABC*^-^ cells*. n = cells, **= p< 0.005*. **(D)** Scanning confocal imaging of mCherry-FRB-MHCKC membrane recruitment and cell shape (brightfield) in AX3 cells. Cells are also expressing an unlabeled membrane-localized FKBP domain (cAR1-2xFKBP, see methods). *t = 00:00 indicates rapamycin addition.* **(E)** Examples (left) and diagram (right) of temporal color projections of cell outlines corresponding to the movie in (D). Blue and yellow times indicate the first and last images in the projection, respectively. **(F)** Average (line) and SEM (shaded area) of cell area before and after MHCKC recruitment by rapamycin addition (dashed line, t = 0). *n = 16 cells*. **(G)** Scanning confocal imaging of RBD and MHCKC membrane recruitment in AX3 cells. *t=00:00 indicates rapamycin addition*. **(H)** Membrane kymograph of Ras activation corresponding to the movie in (G). *Dashed line indicates rapamycin addition, scale bar = 6 minutes*. **(I)** Average (line) and SEM (shaded area) of normalized percentages of the membrane periphery with RBD localization significantly above background before and after MHCKC recruitment by rapamycin addition (dashed line, t = 0). *n = 20 cells*. **(J)** Schematic of the core signaling system controlling cell migration (shaded area) as well as cytoskeletal effectors. Gray arrows indicate previously known interactions while the black arrows indicate the positive feedback from branched actin (**Fig 1A-C, S1A-B**) and the negative feedback from myosin (**Fig 1D-I**) shown here. *Time is in min:sec; scale bars = 5 µm unless otherwise noted*.

We next sought to determine whether cytoskeletal proteins at the cell back also regulate signaling networks. To measure how changing contractility effects signaling, we utilized myosin heavy chain kinase C (MHCKC) to control bipolar thick filament (BTF) assembly of myosin. MHCKC is recruited to the cell cortex after chemoattractant stimulation, leading to myosin phosphorylation and disassembly at specific locations^39,40^. However, since chemoattractant addition triggers multiple events simultaneously, we aimed to develop a method for specifically controlling myosin assembly. To achieve this, we designed a chemically induced dimerization system (CID) utilizing a MHCKC-FRB fusion protein, which is expected to dimerize with cAR1-2xFKBP, a uniformly distributed membrane protein, upon addition of rapamycin (**Fig. 1D-F, Movie 2**).

To confirm BTF disassembly, we engineered the CID system into *myosin II*-null (*mhcA* null) cells “rescued” with GFP-myosin and imaged with total internal reflection fluorescence (TIRF) microscopy. Adding rapamycin led to uniform recruitment of MHCKC to the membrane and an approximately 40% decrease in the intensity of GFP-myosin on the bottom cortex (**Fig. S2A-B**). Imaging cell shape using confocal microscopy revealed that MHCKC recruitment to the cell periphery induced cell spreading, with an approximately 40% increase in cell area, and triggered dynamic changes in cell shape that persisted for at least 20 minutes (**Fig. 1 D-F, Movie 2**).

Increased area and shape change suggested that MHCKC recruitment increased the cells deformability, most likely by reducing its elastic modulus, which is a short time-scale mechanical parameter^41^. This increased deformability then likely makes protrusion formation more likely. To analyze the impact of MHCKC recruitment on STEN activity specifically, we imaged RBD-GFP, an active Ras biosensor as one of the indicators of STEN activation^42^. MHCKC recruitment resulted in a 50% increase in Ras activity at the periphery where protrusions formed (**Fig. 1 G-I, Movie 3**).

Previously, we found that reducing PI(4,5)P2 levels lowers the STEN threshold, increasing the size and number of Ras and PIP3 membrane patches^3^. We sought to determine whether inhibiting myosin would similarly lower this threshold. To address this, we prepared sets of cells having MHCKC recruited to the membrane by pretreatment with rapamycin with untreated cells having no recruitment. We exposed each set to two doses of cAMP (1 nM and 100 nM). At 1 nM cAMP, the pretreated set exhibited a higher average peak response, as measured by depletion of RBD-GFP from the cytoplasm, compared to the untreated set, while at 100 nM cAMP, the two sets responded similarly (**Fig. S2C-D**). Altogether, these findings reveal a negative feedback loop from myosin which increases the STEN activation threshold and inhibits front state formation (**Fig. 1J**).

### Effects of removing branched actin networks on STEN activity

To further investigate negative feedback from the actomyosin network and to probe whether actin populations at the cell front can tune signaling, we inhibited branched actin nucleation with the Arp2/3 inhibitor CK666^43^ and examined changes in actin network morphology and STEN activation. LimE_ΔCoil_, a biosensor for polymerizing actin, localizes to growing pseudopods and macropinosomes in randomly migrating *Dictyostelium* cells^44^. However, within 2 minutes of CK666 treatment, the formation of new pseudopods was significantly inhibited. Interestingly, instead of leaving the cell periphery, LimE redistributed from transient patches and ruffles into a more stable ring-like structure, which began to form after 4 minutes and reached completion after 12 minutes (**Fig. 2A-B, Movie 4**). Because this ring-like actin organization resembled proteins in the actomyosin cortex, we further investigated the effect of CK666 on the localization of Cortexillin I. Cortexillin I is an actin-actin crosslinker, actin-membrane crosslinker, and myosin-binding protein that, as noted earlier, enriches at the cell back, “shadow” waves, and also the cleavage furrow of dividing cells^26,45–49^. Treatment with CK666 led to a substantial increase of approximately 30% in the membrane to cytosol ratio of Cortexillin I relative to DMSO controls (**Fig. S3A-B**). These results show that CK666 treatment not only inhibits branched actin nucleation, but leads to increased “rear cortex-type” assembly.

**Figure 2:**
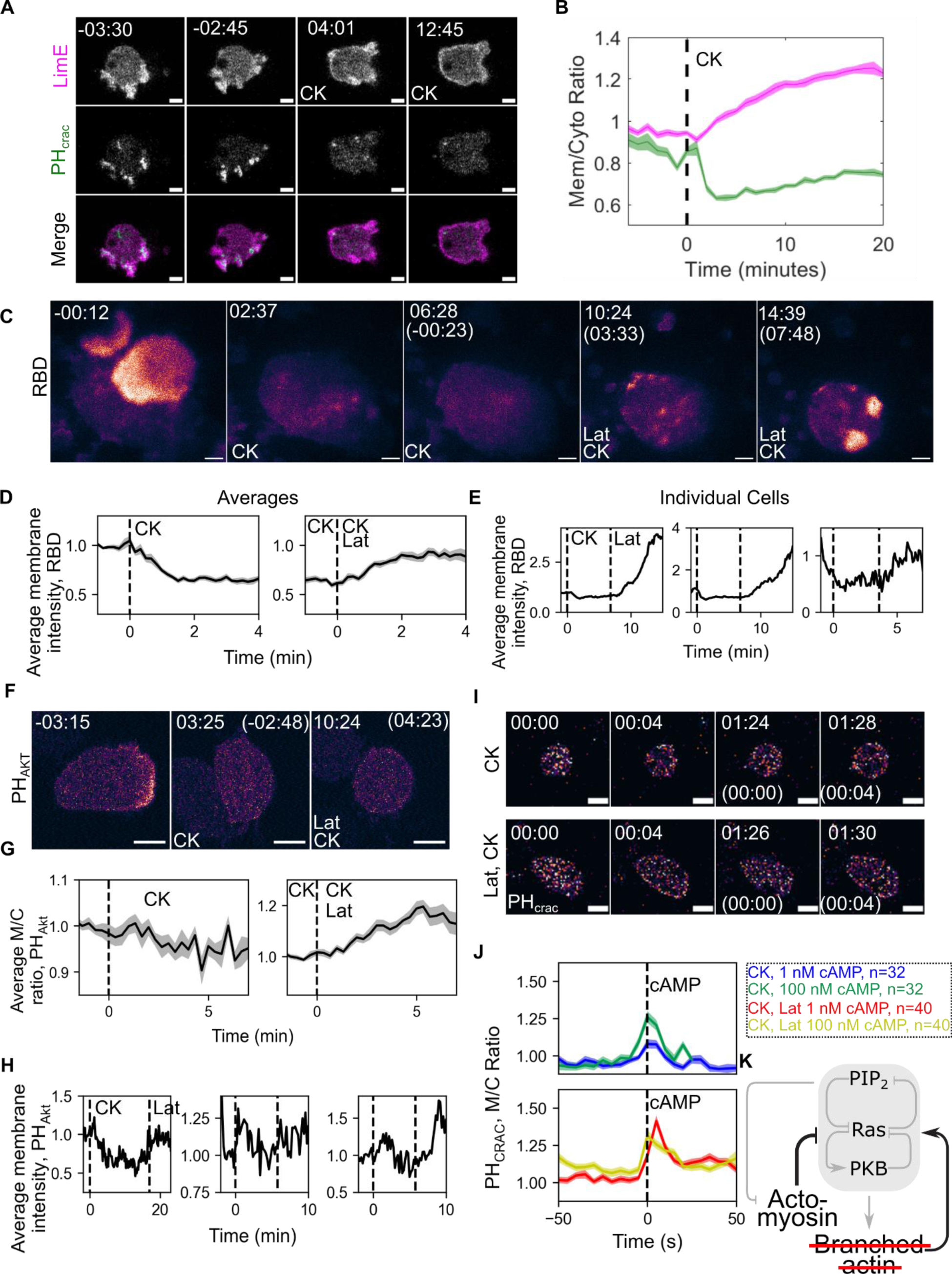
Cortical actin suppresses signaling network activation. **(A)** Scanning confocal imaging of polymerizing actin (LimE_ΔCoil_-RFP) and PIP_3_ levels (PH_CRAC_-YFP) in wild type (AX3) *Dictyostelium* cells before and after treatment with the Arp2/3 inhibitor CK666 (CK). *t = 00:00 indicates CK666 addition*. **(B)** Average (line) and SEM (shaded area) of the membrane-to-cytosol intensity ratio of LimE (magenta) and PH_CRAC_ (green) before and after CK666 addition (dashed line, t = 0). *n = 27 cells*. **(C)** TIRF imaging activated Ras (RBD-mCherry) in electrofused (“giant”) AX3 cells before and after CK666 addition and subsequently latrunculin addition. Cells are incubated in caffeine to raise basal activity levels. *t = 00:00 indicates the addition of CK666 or, in parentheses, latrunculin*. **(D)** Average (line) and SEM (shaded area) of average RBD intensity on the membrane of giant AX3 cells in buffer before and after CK666 addition (left, dashed line, t = 0) or in CK666 before and after latrunculin addition (right, dashed line, t = 0). The left plot excludes data after latrunculin addition while the right excludes data before CK666 addition. *n = 22 cells*. **(E)** Selected individual traces from the dataset in (D). The left plot corresponds to the movie in (C). t = 00:00 indicates CK666 addition; dashed lines correspond to indicated drug addition. **(F)** Scanning confocal imaging of PIP3 levels (RFP-PH_AKT_) in human neutrophil-like (dHL60) cells before and after the addition of CK666 addition and subsequently latrunculin addition. *t = 00:00 indicates the addition of CK666 or, in parentheses, latrunculin*. **(G-H)** Average (G, line), SEM (G, shaded area), and individual traces (H) of the normalized PH_AKT_ membrane-to-cytosol ratio before and after CK666 and subsequently latrunculin addition in dHL60 cells. The right plot in (G) contains both cells starting in buffer and cells pre-incubated in CK666 while the left only contains cells starting in buffer. The left plot in (H) corresponds to the movie in (F), all dashed lines indicate drug addition. *n = 27 cells prior to CK666 addition, 55 cells after CK666 addition and latrunculin addition.* **(I)** Scanning confocal imaging of PIP3 levels in developed AX3 *Dictyostelium* cells treated with CK666 (CK) or CK666 and latrunculin (CK, Lat) before and after the addition of 1 nM cAMP and subsequently 100 nM cAMP*. t = 00:00 indicates the addition of 1 nM cAMP or, in parentheses, 100 nM cAMP.* **(J)** Average (line) and SEM (shaded area) of the membrane-to-cytosol ratio of PH_CRAC_ before and after the indicated dose of cAMP. *n=cells*. **(K)** Arp2/3 experiments reveal negative feedback to signaling from F-Actin left behind after CK666 treatment and positive feedback from branched actin networks. Red line striking out “Branched actin” reflects the CK666 treatment effect. *Time is in min:sec; scale bars = 5 µm*.

To assess the impact of CK666 treatment on signal transduction network (STEN) activity, we employed the biosensor PH_CRAC_, which detects PI(3,4,5)P3^50^. In single cells, PH_CRAC_ overlaps with RBD and LimE in growing pseudopods and macropinosomes. Similar to actin, PIP3 disappeared from these patches following CK666 treatment. Unlike actin, PIP3 did not relocalize elsewhere, suggesting inhibition of the STEN (**Fig. 2A-B**). To further examine how CK666 treatment affects STEN, we measured changes in STEN waves on the bottom cell surface of electrofused giant cells^51^. Initially, these cells displayed propagating waves of RBD along the bottom membrane; however, upon addition of CK666, these waves disappeared within two minutes (**Fig. 2C-E, Movie 5**). These experiments reveal that both PIP3 production, Ras activation, and likely many other STEN activities, are reduced when branched actin is removed and cortical assembly is increased.

To determine whether the decrease in STEN activation was due to the loss of branched actin or the increase in cortical actin, we introduced the actin depolymerizing drug latrunculin to CK666-treated cells to eliminate all remaining actin subpopulations. Significantly, the reduction of F-actin levels in cells lacking branched actin led to the recovery of RBD waves within two minutes (**Fig. 2C-E, Movie 5**). The inhibitory role of cortical actin networks on STEN activity were not limited to *Dictyostelium*. We conducted the same experiment using human neutrophil-like HL60 cells expressing the PIP3 biosensor PH-AKT. Consistent with previous reports, PIP3 was observed in patches at the front of migrating HL60 cells. Upon CK666 addition, these cells rounded up and lost their pseudopods within 2 minutes. Similar to *Dictyostelium*, the elimination of branched actin in neutrophils also resulted in a decrease in PIP3 levels on the membrane. Subsequently, the addition of latrunculin, which removed the remaining actin networks, partially restored PIP3 levels (**Fig. 2F-H, Movie 6**). These findings suggest that the actomyosin cortex suppresses STEN activity and that this phenomenon is evolutionarily conserved.

To test if the inhibition of STEN by the actomyosin cortex was due to an increase in the STEN threshold, we examined the accumulation of PIP3 in response to low (1 nM) and high (100 nM) cAMP stimuli in *Dictyostelium* cells treated with CK666 alone or CK666 and latrunculin in combination. The average peak PIP3 level in response to low cAMP was significantly smaller in cells treated with only CK666 versus cells treated with both latrunculin and CK666. However, there was no significant difference between the two populations when stimulated with high cAMP (**Figs 2I-J, S3**). This relative inhibition indicates that the threshold for STEN was raised, but the system is still functional. Together, these findings suggest that the actomyosin cortex tunes response to receptor inputs by lowering STEN excitability (**Fig. 2K**).

### Myosin II is a critical contributor to negative feedback from the cortex to signaling networks

Because myosin is a critical component of the cortex, we investigated whether removing myosin ameliorates the effect of CK666 on the STEN. CK666 treatment in *myosin*-null cells prevented protrusions and decreased PIP3 levels similar to its effects in wild-type cells. However, unlike the near absence of PIP3 in wild-type cells, *myosin*-null cells treated with CK666 periodically exhibited bright, transient PIP3 patches (**Fig. 3A, Movie S7**). *Myosin*-null cells treated with CK666 had significantly more PIP3 patches than wild type cells (**Fig. 3B**), indicating that myosin is required for the full effect of cortical suppression of STEN. To further elucidate the role of the actomyosin cortex, we applied our method to rapidly dissemble myosin in cells treated with CK666. First, cells were inhibited with CK666, which reduced the number of Ras patches by 95% and immobilized the cell. Then, MHCKC was recruited to the membrane using rapamycin. While disassembling myosin filaments did not restore motility, it caused Ras activation to recover to 40% of its wild type value (**Fig. 3C-E, Movie S8**). Taken together, these results show that myosin is major contributor to cortical feedback on the STEN (**Fig. 3F**).

**Figure 3:**
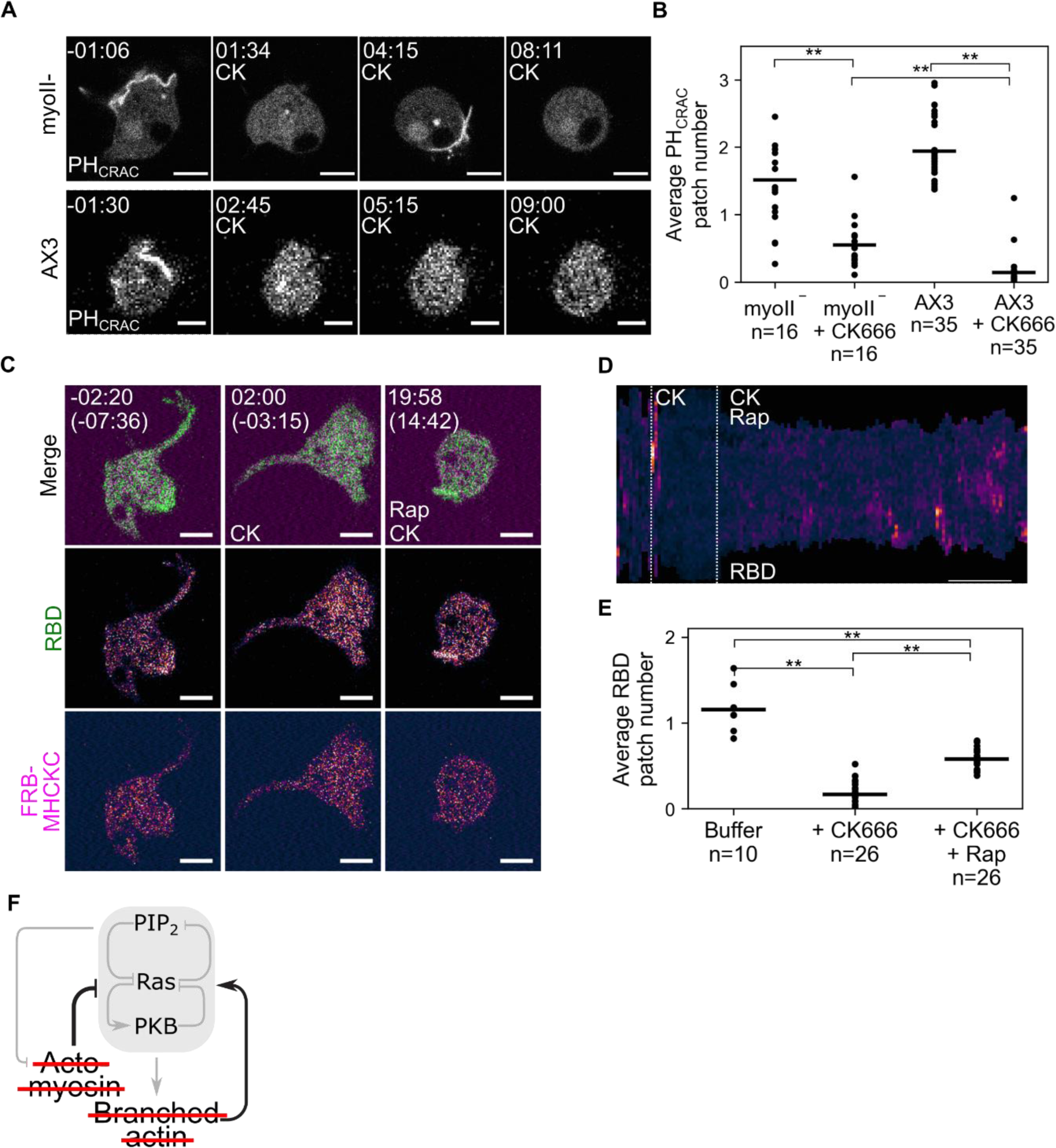
The actomyosin cortex exhibits negative feedback onto signaling networks. **(A)** Scanning confocal images of PIP_3_ levels (PH_CRAC_-YFP) in wild type (AX3) and *myosin II*-null (*myoII-*) *Dictyostelium* cells before and after CK666 addition. *t = 00:00 indicates CK666 addition.* **(B)** Average (lines) and individual (dots) mean number of PIP_3_ patches over time before and after CK666 addition in AX3 and *myoII^-^* cells. *n = cells, ** = p < 0.005.* **(C)** Scanning confocal images of Ras activation (RBD-EGFP) before and after CK666 addition and subsequently mCherry-FRB-MHCKC membrane recruitment in AX3 cells. Cells are also expressing an unlabeled membrane-localized FKBP domain (cAR1-2xFKBP). *t = 00:00 indicates the addition of CK666 or, in parentheses, rapamycin*. **(D)** Membrane kymograph of Ras activation corresponding to the movie in (C). *Dashed lines represent the addition of specified drug, scale bar = 5 minutes*. **(E)** Average (lines) and individual (dots) mean number of RBD patches over time before and after CK666 addition and subsequently MHCKC recruitment by rapamycin addition. *n = cells, ** = p < 0.005*. **(F)** The partial reversal of CK666’s effect on signaling by ablating myosin (**Fig 3A-E**) indicates that the actomyosin cortex as an ensemble feeds back negatively on cell signaling. Red lines indicate that Arp 2/3 and Myosin have been eliminated or reduced. *Time is in min:sec; scale bars = 5 µm unless otherwise noted*.

### Using RacE as a tool to probe the role of cortical actin in STEN regulation

To further probe the role of the actomyosin cortex in signaling, we turned to RacE, a member of the Rac/Rho subfamily which has been shown to regulate linear actin nucleation, actin crosslinking, Myosin contraction, and cortical viscoelasticity and tension^52–56^. To abruptly increase RacE activity in migrating cells and assess signaling impacts, we designed a CID system to recruit the RacE GEF, GXCT^57^, to the cell membrane. Recruiting the GEF domain of GXCT to the cell membrane by adding rapamycin led to a phenotype with cells dramatically flattening and undulating rapidly around the perimeter, while losing well-defined protrusions and motility (**Fig. 4A-D, Movie S9**). LimE localization shifted from a few broad patches to a uniform distribution along the cell perimeter. Quantitative analysis showed a 40% increase in F-actin cortical localization within 8 minutes after rapamycin addition (**Fig. 4E-G, Movie S10**). Similarly, the amount of actin crosslinker dynacortin increased significantly upon GXCT recruitment, consistent with the role of RacE in the recruitment this cytoskeletal protein to the cortex^52^(**Fig. S4A-C**). Furthermore, we confirmed that the changes in actin were not due to an unknown role of RacE in activating Arp2/3 by repeating the experiment in cells treated with CK666. When we recruited GXCT to the membrane, there was still a significant increase in actin polymerization, indicating that RacE activates actin polymerization independently of Arp2/3 (**Fig. S4D-E**).

**Figure 4:**
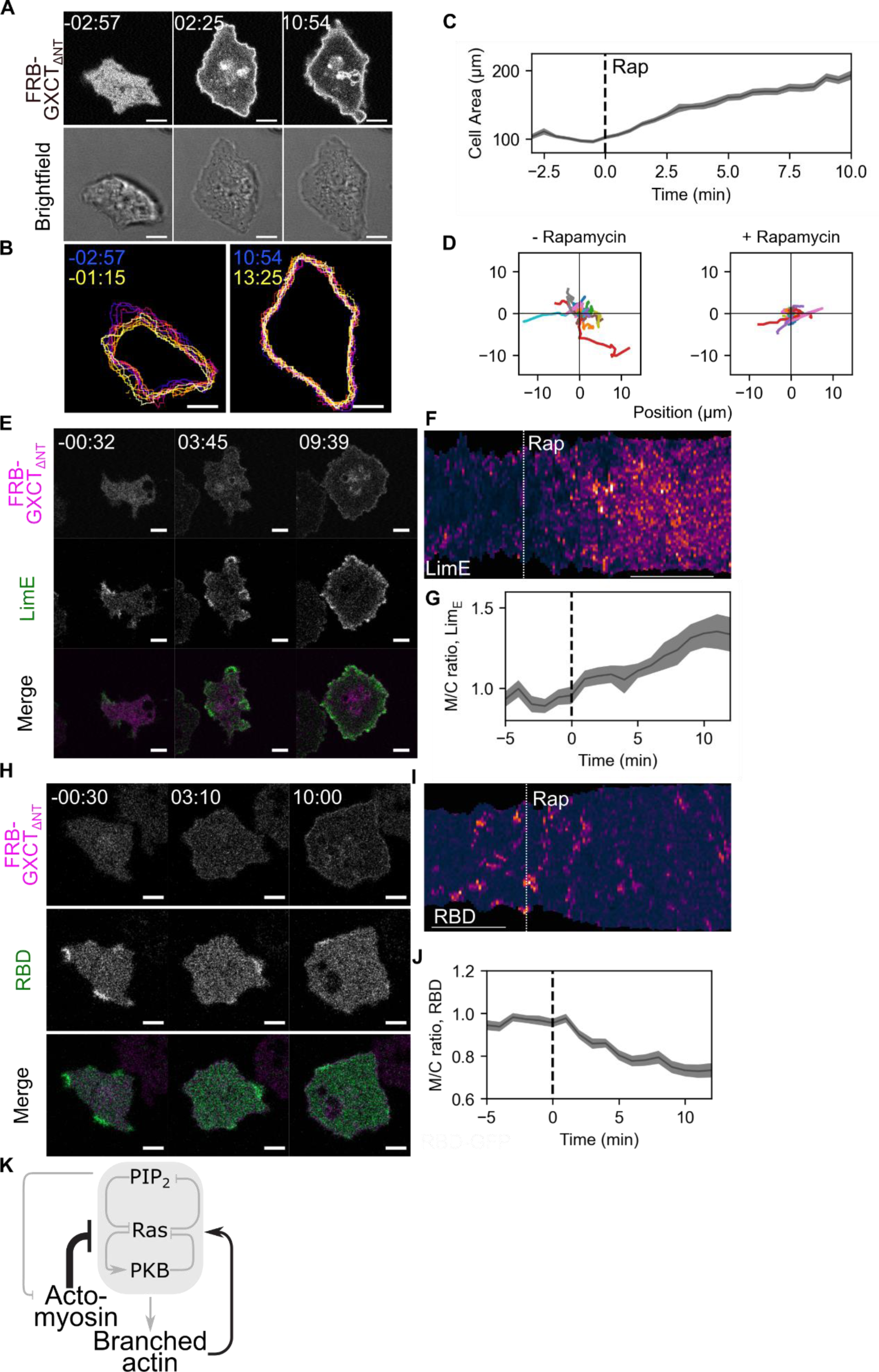
Increasing the abundance of the actomyosin cortex using RacE leads to signaling inhibition. **(A)** Scanning confocal imaging of RacE-GEF (mCherry-FRB-GXCT_ΔNT_) membrane recruitment and cell shape in wild type (AX3) *Dictyostelium* cells. Cells are also expressing an unlabeled membrane-localized FKBP domain (cAR1-2xFKBP). *t = 00:00 indicates rapamycin addition.* **(B)** Temporal color projections of cell outlines corresponding to the movie in (A). Blue and yellow times indicate the first and last images in the projection, respectively. **(C)** Average (line) and SEM (shaded area) of cell area in AX3 cells before and after GXCT recruitment (dashed line, t=0). *n = 35 cells.* **(D)** Traces of cell movement in AX3 cells 200 seconds before and after GXCT recruitment. *n = 35 cells.* **(E)** Scanning Confocal imaging of GXCT recruitment and polymerizing actin (LimE_ΔCoil_-EGFP) in AX3 cells. *t = 00:00 indicates rapamycin addition.* **(F)** Membrane kymograph of LimE from the movie in (E). *Dashed line indicates rapamycin addition, scale bar = 5 minutes.* **(G)** Average (line) and SEM (shaded area) membrane-to-cytosol ratio of LimE before and after GXCT recruitment by rapamycin addition (dashed line, t = 0). *n = 8 cells.* **(H)** Scanning Confocal imaging of GXCT recruitment and Ras activation (RBD-EGFP) in AX3 cells. *t = 00:00 indicates rapamycin addition.* **(I)** Membrane kymograph of RBD from the movie in (H). *Dashed line indicates rapamycin addition, scale bar = 5 minutes.* **(J)** Average (line) and SEM (shaded area) membrane-to-cytosol ratio of RBD before and after GXCT recruitment by rapamycin addition (dashed line, t = 0). *n = 9 cells. Time is in min:sec.* **(K)** Increasing the abundance of the actomyosin cortex (thick black arrow) inhibits the core signaling module. *Scale bars = 5 µm unless otherwise noted*.

To determine the effects of RacE activation on STEN, we observed changes in the localization of the active Ras biosensor, RBD, after GXCT recruitment. Before recruitment, cells displayed several large, intense, and long-lived RBD patches along their periphery. However, upon activating RacE, the RBD patches became smaller, weaker, and shorter-lived, with a potential increase in the total number of patches. The amount of Ras activation on the cell periphery dropped by 30% within 8 minutes (**Fig. 4H-J, Movie S11**). Taken together, these results suggest that RacE activation is able to increase actin polymerization in an Arp2/3-independent manner and thereby inhibit STEN activation (**Fig. 4K**).

### RacE reversibly and locally inhibits STEN and triggers arp2/3-independent protrusions

To reversibly and locally control RacE activity, we designed an iLID-based optogenetic system^58^ to recruit GXCT to the membrane by applying blue light to cells. In this system, the GEF domain of GXCT was fused to a SSPB domain which binds to a membrane-localized iLid domain (see methods) when exposed to blue light. Similar to the CID system, uniform light stimuli led to rapid cell flattening and cessation of movement, which rapidly reversed upon removal of illumination (**Fig. 5A-C, Movie S12**). We then examined how transiently increasing RacE activity affected propagating basal RBD waves. Recruiting GXCT caused a sudden decrease in RBD wave activity, which was quickly reversed upon removal of blue light and dissociation of GXCT (**Fig. 5D-F, Movie S13**).

**Figure 5:**
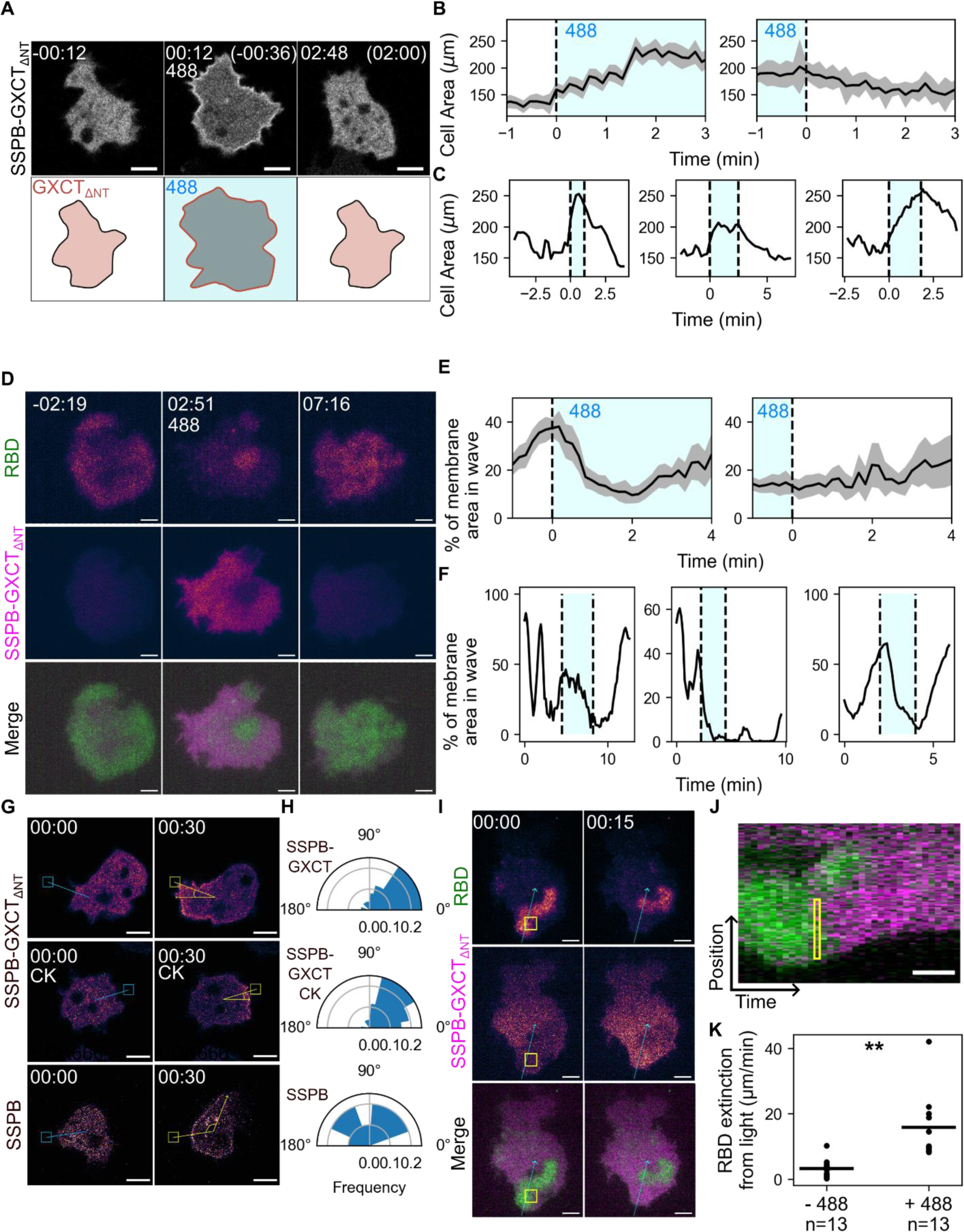
RacE reversibly and locally control cell signaling and actin polymerization. **(A)** Scanning confocal imaging of RacE-GEF (tagRFP-SSPB-GXCT_ΔNT_) optical membrane recruitment in wild type (AX3) *Dictyostelium cells*. Cells are also expressing an unlabeled membrane-localized iLID domain (N150-ILID, see methods). *t = 00:00 indicates blue light exposure or blue light loss (parentheses)*. **(B)** Average (lines) and SEM (shaded area) of cell area before and after GXCT membrane recruitment (left, dashed line) or GXCT membrane dissociation (right, dashed line). *n = 17 cells (left) and 6 cells (right)*. **(C)** Individual traces of cell area in cells before, during (shaded area), and after GXCT recruitment. The plot on the left corresponds to the movie in (A). **(D)** TIRF imaging of Ras activation (RBD-emiRFP670) before, during, and after GXCT recruitment in electrofused (“giant”) AX3 cells. Cells are treated with 50 µg/ml Biliverdin to activate emiRFP670 fluorescence. *t = 00:00 indicates blue light exposure.* **(E)** Average (lines) and SEM (shaded area) of the percentage of the cell membrane with RBD localization significantly above background before and after GXCT membrane recruitment (left, dashed line) or GXCT membrane dissociation (right, dashed line). *n = 7 cells*. **(F)** Individual traces of the percentage of the cell membrane with RBD localization significantly above background in giant cells before, during (shaded area), and after GXCT recruitment. The plot on the left corresponds to the movie in (D). **(G)** Scanning confocal imaging of cell protrusion formation after local recruitment of SSPB-GXCT or SSPB alone. +CK indicates cells were pre-treated with CK666. Boxes indicate the region of blue light exposure; arrows form a line between the center of the protrusion formed after stimulation and the center of the cell. *t = 00:00 indicates the last timepoint before blue light exposure.* **(H)** Angular histograms of the angle formed between the center of the protrusion and the location of GXCT recruitment relative to the cell center as demonstrated in (G). *n = 26 cells, SSPB-GXCT and SSPB-GXCT + CK666. n = 15 cells, SSPB.* **(I)** TIRF imaging of RBD membrane localization before and after local GXCT recruitment in giant AX3 cells. The yellow box indicates the region of blue light exposure and the blue arrow indicates the line and direction for linear kymograph creation. *t = 00:00 indicates the last timepoint before blue light exposure.* **(J)** Linear kymograph of GXCT recruitment and RBD intensity corresponding to the blue line in (I). Yellow box indicates the approximate region of blue light exposure. *Scale bar indicates 20 seconds*. **(K)** Quantification of the motion of RBD waves away from the region of GXCT recruitment (see methods). Because this measurement is a composite of RBD translocation and disappearance it is not a true velocity. *n = cells, ** = p < 0.005. Time is in min:sec; scale bars = 5 µm unless otherwise noted*.

Next, we took advantage of our optogenetic system to assess the morphological and signaling effects of local increases in RacE activity. Interestingly, despite the ability of RacE to turn off Ras activity, the localized recruitment of GXCT induced the formation of protrusions expanding from the recruitment site towards the source of the blue light. These protrusions were observed to form even in the presence of CK666, suggesting that RacE-triggered actin polymerization can occur independently of Arp 2/3 (**Fig. 5G-H, Movie S14**). However, the lifetime and length of these protrusions is lower than typical chemoattractant-induced pseudopods. Moreover, recruiting GXCT to specific regions on the basal surface of the cell led to the local extinguishment or redirection of propagating RBD waves (**Fig. 5I, Movie S15**)

### Feedback loops from the cytoskeleton can generate cell polarity

Our results suggest the presence of two different cytoskeletal feedback loops. To determine how these loops could affect excitability and cell polarity, we turned to a computational model. The core of the model is an activator-inhibitor system (**Fig. 6A**) in which the activator positive feedback is implemented as double negative feedback between Ras and anionic lipids, including PIP2, and the slow inhibition is between Ras and PKB^24^. Simulations of this core system show occasional firings randomly distributed around a cell perimeter (**Fig. 6B**). These elevated regions of activity propagate for several seconds before they extinguish. The complementary pattern between Ras and PIP2 was used as a readout.

**Figure 6:**
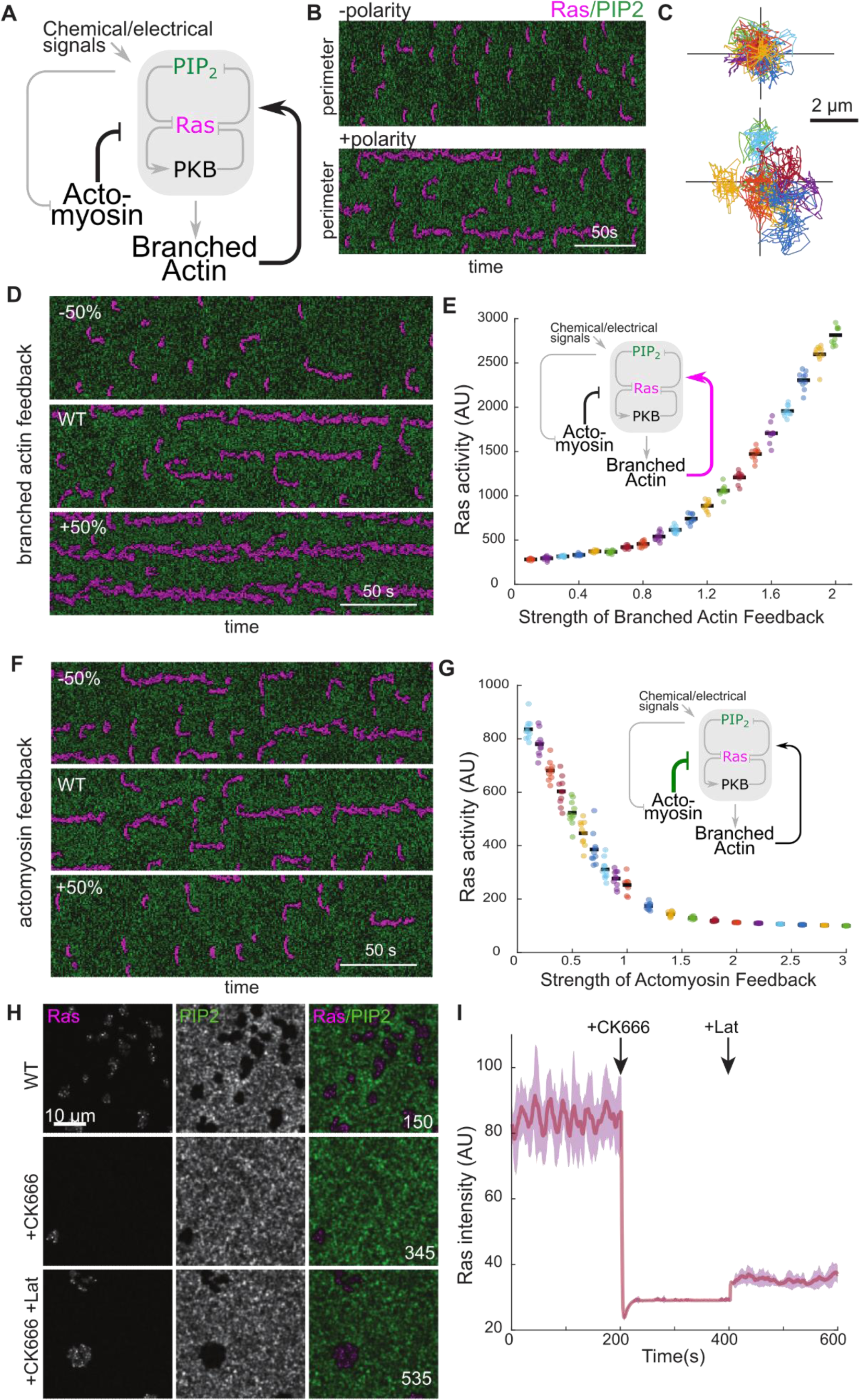
Cytoskeletal networks can create polarity by feeding back onto a core signaling module. **(A)** Schematic showing the signal transduction network involving Ras/PIP2 and PKB and how they couple to the two types of actin feedback loops. **(B)** Kymographs showing activity around the cell perimeter as function of time. PIP2 and Ras are shown in green and magenta respectively. The top and bottom correspond to simulations without and with the feedback loops, respectively. **(C)** Simulated cell trajectories of 10 cells each with feedback loops turned off (top) and on (bottom). **(D)** Kymographs showing wave activity across cell perimeter for varying strengths of the branched actin feedback. **(E)** Total Ras activity around the cell perimeter with respect to the strength of the branched actin feedback. Wildtype corresponds to a strength of 1. Black bar denotes the mean of 10 simulations per strength. Simulations with total Ras activity less than 500 showed no firings. **(F)** Kymographs showing wave activity across cell perimeter for varying strengths of the actomyosin feedback. **(G)** Total Ras activity around the cell perimeter with respect to the strength of the actomyosin feedback. Wildtype corresponds to a strength of 0.4. Black bar denotes the mean of 10 simulations per strength. Simulations with total Ras activity less than 200 showed no firings. **(H)** Frames from a 2D simulation of effects of adding CK666 and then Latrunculin to wild type cells. The three rows represent waves in the wildtype cell, waves after CK666 addition, and subsequent Latrunculin treatment. **(I)** The total Ras activity for simulations as in panel H. The solid line and the shaded area represent the mean ± 1 standard deviation. In all the simulations the CK666 effect is incorporated at the 200 s and the additional Latrunculin effect is added at the 400 s. *Scale bars = 10 microns*

To the basic network, we add two complementary feedback loops (Black lines in **Fig. 6A**). The first loop implements the feedback between branched actin and Ras at the cell front while the second implements feedback between actomyosin and Ras at the rear. Details of the mathematical model are given in the Methods. Simulations of the system with added cytoskeletal feedback have the same complementarity as before, but Ras shows increased persistence, characterized by localized long-lived regions of elevated Ras with minimal lateral propagation (**Fig. 6B**).

To illustrate the effect of these feedback loops on cell movement, we first plotted the trajectories of simulated cells (see Methods) migrating randomly in the absence of branched actin and actomyosin feedback (**Fig. 6C, top**) and compared it to those in the presence of these loops (**Fig. 6C, bottom**). Cells without feedback remained near their starting point, while the addition of feedback loops enabled cells to move farther away.

To characterize the effect of the two loops separately, we altered the strength of each loop individually while holding the other constant and quantified the total Ras activity. Increases in branched actin feedback led to elevated Ras-GTP levels, greater persistence, and wider patches (**Fig. 6D, E**). Increases in actomyosin feedback had the opposite effect (**Fig. 6F, G**).

Finally, we carried out simulations in a two-dimensional surface to recapitulate basal waves and recreated the experiments of **Fig. 2C-D**. Initially, waves are seen, but after simulating the effect of CK666 addition through the removal of the branched actin feedback, they disappear. The subsequent removal of the actomyosin feedback loop, mimicking Latrunculin addition, leads to a partial recovery of the Ras waves (**Fig. 6H; Movie S16**). Quantitating the effect over multiple simulations shows that total Ras activity decreases greatly when CK666 is added and partial recovery upon Latrunculin treatment (**Fig. 6I**).

## Discussion

Much previous work has focused on control of the cytoskeleton by Ras/PI3K signal transduction networks, particularly in the context of chemotaxis, and many studies have focused on the reaction of cytoskeletal elements to mechanical forces, but feedback from the cytoskeleton to “upstream” signal transduction events has only occasionally been considered. Using synthetic biological techniques, we have uncovered two crucial feedback mechanisms from different actin networks that characterize the front and back regions of the cell cortex. A front-promoting mechanism is supported by the observation that increases in Arp2/3 actin strongly boost Ras/PI3K activation, while reduction of F-actin decreases the activation. A back-promoting mechanism is indicated by multiple observations. First, acute reduction of myosin assembly enhances Ras/PI3K activation and heightens sensitivity to chemotactic stimuli, indicating that the actomyosin cortex suppresses Ras activation. Second, selective inhibition of branched actin nucleation nearly eliminates Ras activity; this inhibition is mitigated by depolymerizing the bulk of actin and decreasing myosin assembly, as well as in cells lacking myosin. Collectively, these observations indicate that the inhibition depends on the cortical actomyosin network. Third, recruitment of RacE GEF, which promotes actin crosslinking and formin nucleation, substantially reduces Ras/PI3K activation and yet can still induce protrusions. This indicates that cells can suppress Ras/PI3K signaling locally or globally by increasing the abundance of the actomyosin cortex.

When taken together with interactions discovered previously, the mechanisms we have delineated create two positive feedback loops between signal transduction and cytoskeletal networks. A front-promoting loop becomes clear by merging earlier findings that Ras/PI3K activation promotes Arp2/3 nucleation with our observations here that branched actin activates these signal transduction networks^59^. A back-promoting loop emerges by bringing together our studies here that myosin assembly and strengthening the actomyosin cortex inhibits Ras/PI3K activation with prior work that Ras/PI3K activation leads to activation of myosin heavy chain kinase and consequent disassembly of myosin^12,36^. In human neutrophil-like cells, Ras has recently been shown to reverse back polarity through its downstream target AKT1, indicating a similar relationship may exist^60^. The existence of these front- and back-promoting loops points to reciprocal interactions between Ras/PI3K signal transduction and the branched actin and actomyosin cytoskeletal networks. These interconnected networks provide a molecular scheme for generating cell polarity and for integrating the chemical and mechanical cues that a cell would encounter in navigating a complex environment

Previous investigators have speculated that polarity is established by dual front and back positive feedback loops, but these findings differ from ours in several ways. First, much of the work on these feedback loops focuses on direct feedback between different cytoskeletal regulators such as Rac and Rho and do not consider a role for upstream signaling events like Ras/PI3K activation. Second, while previous models have also described positive feedback to PIP3 from Rac^8,11,15–18^, we show here that the actomyosin cortex is a direct negative regulator of Ras/PI3K signaling, even in cells without Arp2/3 activity. These findings are not inconsistent, but our work reveals an additional critical layer of regulation.

The mechanism of negative feedback from the actomyosin cortex to signaling is not yet clear, but there are several attractive possibilities. One possibility is that integrity of the underlying cortex influences the fluidity of membrane. Lower fluidity in regions with high cortical density may slow the diffusion of signaling activators, locally lowering excitability. Supporting this, multiple studies have reported lower membrane fluidity in back regions where actomyosin density should be high^61–63^. Another possibility is related to the finding that anionic lipids such as PI(4,5)P2 on the inner leaflet of the membrane inhibits Ras/PI3K signaling^6^. Electrostatic interactions between molecules in the actomyosin cortex and the lipids, such as through cortexillin and PI(4,5)P2^64^, or the enzymes that regulate them may contribute to the enrichment of these inhibitory lipids at the cell back. Recent work has indicated that actin and the membrane are more tightly associated at the cell back, indicating a stronger interaction between the membrane and the actomyosin cortex than branched actin^63^. Finally, force generation by the actomyosin cortex could alter membrane tension or curvature, leading to the recruitment or activation of negative regulators specifically at the cell back^65–69^.

A cell migrating though its natural environment towards a chemical cue encounters a complex mix of other cells, the extracellular matrix, and shear forces. These physical features are usually integrated with chemical cues to determine the optimal path towards a destination. One might imagine that the cell is like a self-driving car, following a GPS signal to a destination: if the car merely followed a pre-planned route, it would soon collide with an unmarked obstacle. To safely reach its goal, the car must use sensors to detect other cars or road work. While cells have well-characterized biochemical sensors, their methods for detecting mechanical stimuli are less well-understood, nor is it clear how cells integrate these two cues. Recent studies have suggested various physical properties, like nuclear stiffness and membrane curvature, may help steer cells in crowded environments^69–71^. Our findings, combined with long-standing evidence that physical forces alter the properties of the cytoskeleton, suggest an attractive hypothesis. These physical cues could locally alter the balance between actomyosin and branched actin. As we have shown here, altering this balance has quite significant effects on the same signaling nodes that are also controlled by chemical stimuli. Therefore, the core excitable system we and others have described likely serves as an integrator of diverse environmental signals to bring about successful navigation^72^.

Our results also have important implications for current and future therapeutic interventions. Currently, many cancer therapies target specific signaling molecules involved in cell growth and migration. Layers of redundancy and feedback from the local environment may compensate for inhibition at any single node. Targeting the cytoskeleton in combination with molecules like Ras and PI3K may be critical for preventing tumor growth and metastasis^73–75^.

## Methods

### Cells and plasmids

*Dictyostelium discoideum* cells were cultured in HL5 media^76^ for a maximum of 2 months after thawing from frozen stock. AX3 cells were obtained from the R. Kay laboratory (MRC Laboratory of Molecular Biology, UK), *abnABC^-^* cells were generated with homologous recombination previously in the Devreotes Lab. *mhcA*^-^ cells were obtained from the D. Robinson Lab (JHU). Female human neutrophil-like HL-60 cells stably expressing RFP-PH_AKT_ was previously created in the Devreotes lab^77^. Cells were cultured in RPMI medium 1640 with L-glutamine and 25 mM HEPES (Gibco; 22400-089) supplemented with 15% heat-inactivated fetal bovine serum (FBS; Thermo Fisher Scientific; 16140071) and 1% penicillin-streptomycin (Thermo Fisher Scientific; 15140122). Cells were split at a density of 0.15 million cells/ml and every 3 days. 1.3% DMSO was added to cells at a density of 0.15 million cells/ml to trigger differentiation 5-7 days before imaging.

Plasmids were introduced into *Dictyostelium* cells using electroporation^78^. To improve efficiency, heat-killed *Klebsiella aerogenes* was added after transformation. In chemically-induced dimerization (CID) experiments (**Fig. 1D-I**, **Fig. 3C-E**, **Fig. 4, Fig S4**), all cells were transformed with an unlabeled cAR1-tandem FKBP (cAR1-FKBP-FKBP) in a pCV5 vector as well as an FRB-tagged protein in a separate pCV5 vector. cAR1 was selected because it is a uniform membrane protein^79^ and will therefore recruit FRB-tagged proteins to the membrane evenly upon rapamycin addition (**Fig. 1D**). mCherry-FRB-MHCKC was created using standard restriction digest cloning starting with a GFP-MHCKC construct (from the D. Robinson lab). mCherry-FRB-GXCT_ΔNT_ was made by inserting the PH and RhoGEF (DH) domains of GXCT ^80^ from a GFP-GXCT construct (gift from the M. Iijima lab, JHU) into a pCV5 plasmid containing mCherry and FRB. While both cAR1-2xFKBP and FRB vectors contain G418-resistance markers, a significant population of cells simultaneously transformed with both have significant FRB protein expression and membrane recruitment.

For optogenetic experiments (**Fig. 5**), cells were transformed with hygromycin-resistant vector pDM358^81^, containing an iLID^58^ domain linked to the myristoylation domain of PKBR1 (n150-ILID), which was previously shown to localize to the membrane ^3^. Cells were subsequently transformed with SSPB(R73Q)-tagRFP-GXCT_ΔNT_ in a pCV5 vector (**Fig. 5A-C**, **Fig. 5G-H**) created by In-Fusion cloning or a dual-expression G418 resistance vector (O1N), containing mCherry-SSPB(R73Q)-GXCT_ΔNT_ and RBD-emiRFP670 (**Fig. 5D-F**, **Fig. 5I-K**) created by GoldenBraid cloning^82^.

pDM358 RBD-EGFP (**Fig. 1A**, **Fig. 1G-I**, **Fig. 3C-E**, **Fig. 4H-J**), LimE_ΔCoil_-EGFP (**Fig. 4E-H, S1A-C**), pDM358 PH_CRAC_-YFP/LimE_ΔCoil_-RFP (**Fig. 2A-B**, **Fig. 2 I-J**), and pDM358 PH_CRAC_-YFP (**Fig. 3A-B**) were previously created in the Devreotes lab^3^. Hygromycin resistant vector pDRH EGFP-MyosinII (**Fig. S2A-B**), pDRH-Cortexillin I (**Fig. S3A-B**), and pDRH-Dynacortin (**Fig. S4D-E**) were created in the the D. Robinson lab. pDM358 EGFP-HSPC300 (**Fig. S1**) was a gift from the C. Huang lab (JHU).

### Microscopy

For *Dictyostelium* imaging experiments, cells were seeded in 8-well Lab-Tek chambers (Thermo-Fisher, 155409) and left to settle for ten minutes before the media was gently aspirated and replaced with Development Buffer (DB) (5 mM NA_2_HPO_4_, 5 mM KH_2_PO_4_, 1mM CaCl_2_, 2 mM MgCl_2_). In experiments with non-electrofused vegetative cells, cells were allowed to sit for an hour starving in DB prior to imaging to decrease photosensitivity. For experiments requiring cAMP stimulation (**Fig. S2C-D, Fig. 2I-J**), growth-phase cells were washed and suspended in DB at a density of 2 × 10^7^ cells/ml. Cells were then developed by being shaken for 1 hour and subsequently pulsed with 50–100 nM cAMP every 6 minutes and shaken for 4 hours.

For dHL60 cells, cells were seeded in Lab-Tek 8-well chambers coated with fibronectin at approximately 35 µg/cm and allowed to adhere for one hour prior to imaging. Directly prior to imaging, cells were treated with 200 nM N-Formyl-Met-Leu-Phe (FMLP) to encourage migration.

Laser scanning confocal imaging was carried out on two microscopes: a Zeiss AxioObserver inverted microscope with an LSM800 confocal module and a Zeiss AxioObserver with 880-Quasar confocal module & Airyscan FAST module. On the LSM800 microscope, GFP and YFP proteins were excited with a solid-state 488 nm laser and on the 880 with an argon laser. mCherry and other red fluorescent proteins were excited with a solid-state 561 nm laser on both systems. Emission wavelengths collected were chosen to avoid overlap between GFP and mCherry emission profiles. All imaging was done with 63X/1.4 PlanApo oil DIC objectives and appropriate raster zoom. Brightfield images acquired using a transmitted-photomultiplier tube (T-PMT) detector.

For Total Internal Reflection Fluorescence (TIRF) imaging, experiments measuring the response of RBD to CK666 and latrunculin (**Fig. 2C-E**) were performed on a Nikon TiE microscope with a solid state 561 nM laser for mCherry excitation, a 100x/1.49 Apo TIRF objective, an RFP emission filter set, and a Photometrics Evolve 512 EMCCD camera. Experiments measuring the response of RBD to RacE GEF recruitment (**Fig. 5D-F, Fig. I-K**) were performed on a Nikon Ti2-E microscope with an iLas2 Ring-TIRF and optogenetics module (GATACA Systems). Red fluorescent proteins were excited with a 561 nM solid-state laser and far-red proteins were excited with a 647 nm solid-state laser. Global and local recruitment were accomplished using a 488 nM solid-state laser connected to the iLas2 system. Cells were imaged using a 60x/1.49 Apo TIRF objective and a Hamamatsu FusionBT camera.

### Cell fusion

Protocol adapted from (Miao et al., 2019).Growth-phase cells were washed twice with and then resuspended in SB (17 mM Sorensen Buffer containing 15 mM KH_2_PO_4_ and 2 mM Na_2_HPO_4_, pH 6.0) at a density of 1.5 × 10^7^ cells/ml. 3 ml of cells were put into a 15-ml conical tube and rolled gently for ∼ 30 min, to promote cell clustering. 800 μl of rolled cells were transferred to a 4-mm-gap electroporation cuvette (Bio-Rad, 1652081), using pipette tips whose edges were cut off to avoid breaking clusters. Cells were then electroporated using the following settings: 1,000 V, 1 μF three times, with 1–2 s between pulses. Then, 30 μl of cells was transferred to the center of a well in an 8-well chamber and was left still for 5 min. 370 (**Fig. 2C-E**) or 470 (**Fig. 5D-F**, **Fig. 5I-J**) μl of SB containing 2 mM CaCl_2_ and 2 mM MgCl_2_ was added to the well and was pipetted briefly to suspend the cells evenly. After allowing cells to settle for 10 minutes, all media together with excess floating cells were removed and 450 μl of new SB plus 2 mM CaCl_2_ and 2 mM MgCl_2_ was gently added to the well against the wall. Cells were then allowed to recover for 1 h before imaging. 50 µg/ml Biliverdin was added to cells expressing RBD-emiRFP670 (**Fig. 5D-F**, **Fig. 5I-J**) during the rolling stage to activate emiRFP970 fluorescence.

### Preparation of Reagents and inhibitors

Rapamycin (Millipore Sigma, 553210) was dissolved in DMSO to a concentration of 10 mM. Then, 1 µl aliquots were diluted 1:200 in the imaging buffer of the experiment (DB or SB plus 2 mM CaCl_2_ and 2 mM MgCl_2_) to a 10x concentration (50 µM). CK666 (Millipore Sigma, 182515) was dissolved in DMSO to concentration of 100 mM. Then, 0.5 µl aliquots were diluted 1:100 in warm imaging buffer of the experiment (DB, SB plus 2 mM CaCl_2_ and 2 mM MgCl_2,_ or RPMI) to a 10x concentration (1 mM). 2.5 µl aliquots of 1 mM latrunculin in DMSO (Millipore Sigma, 428026) were diluted 1:20 in the imaging buffer of the experiment, creating the 50 µM 10x solution used for the experiments (DB, SB plus 2 mM CaCl_2_ and 2 mM MgCl_2,_ or RPMI). cAMP (Millipore-Sigma, A6885) was dissolved to 1 mM in water and then diluted to a 10x working concentration (1 µM or 10 nM) in DB. Caffeine (Millipore-Sigma, 1085003) was dissolved to 1 M in water and then diluted to a 10x concentration (30 µM) in SB plus 2 mM CaCl_2_ and 2 mM MgCl_2_. FMLP was dissolved in DMSO to a concentration of 10 mM and then individual aliquots were diluted in RPMI to a 10X concentration (2 µM).

### Chemically induced dimerization and inhibitor experiments

When imaging FRB protein recruitment without any additional inhibitors (**Fig. 1D-I**, **Fig. 4, Fig. S4**), 50 µl of 50 µM rapamycin was added to a well containing 450 µl of buffer at the indicated time for a final concentration of 5 µM. Similarly, in experiments with CK666 alone (**Fig. 2A-B, Fig. S3, Fig. 3A-B**), 50 µl of 1 mM CK666 was added to a well containing 450 µl of buffer at the indicated time for a final concentration of 100 µM. For experiments with rapamycin and CK666 (**Fig. 3C-E, Fig. S4A-C**) or CK666 and latrunculin (**Fig. 2C-H**), 50 µl of each of the drugs was added to a well containing 400 µl of buffer at the indicated times for a final concentration of 5 µM Rapamycin, 100 µM CK666, and 5 µM latrunculin. In **Fig. 2C-E**, cells were pre-incubated with 3 mM caffeine, and all drugs were diluted in buffer containing 3 mM Caffeine.

For cAMP stimulation experiments, cells were pre-treated in 400 µl DB containing rapamycin (**Fig. S2C-D**), CK666, or CK666 and latrunculin at the concentrations indicated above for 30 minutes. Then, 50 µl of 10 nM cAMP was added to the well during imaging followed at least a minute later by 50 µl of 1 µM cAMP for final concentrations of 1 nM and 100 nM respectively.

In **Fig. S2A-B**, the + rap population was treated with 5 µM rapamycin at least 30 minutes prior to imaging. For cells pre-treated with CK666 in **Fig. 2F-G** and **Fig. 3C-E**, cells were treated with 100 µM CK666 at least 30 minutes prior to imaging.

### Optogenetics

In **Fig. 5A-C** and **5G-H**, experiments were performed on the Zeiss 800 LSM. For global activation (**Fig. 5A-C**), the 488 laser was switched on and scanned across the entire field of imaging for the indicated time. For local activation (**Fig. 5G-H**), the microscope’s bleaching tool was used to scan the 488 laser in an approximately 1 µm square 1 µm away from the cell edge.

In **Fig. 5D-F** and **5I-K**, experiments were performed on the iLas TIRF system. For global activation (**Fig. 5D-F**), the cells were exposed to 488 laser light at every timepoint for the indicated time. For local activation (**Fig. 5I-K**), the microscope’s targeted illumination tool was used to scan the 488 laser in an approximately 300 µm square on the cell membrane at the indicated location.

### Image analysis

To quantify biosensor intensity on the membrane in confocal images, single cells were selected for analysis using cell health, biosensor expression, and FRB/SSPB protein recruitment (if applicable). Cells were then tracked manually using the Manual Tracking plugin in FIJI^83^. After manual tracking, masks of individual cells were created in MATLAB (Mathworks). Background was subtracted from the image by applying a 2D gaussian blur filter with a high σ value (100) to the original image and then subtracting the resulting image from the original. Then, a threshold intensity value was calculated on the background-subtracted image using Otsu’s method^84^ and applied to a smoothed version of the background-subtracted image created using a 2D gaussian blur filter with a low σ value (2). Masks were then modified by performing a morphological closing operation followed by a hole-filling operation. The selected cell was then identified by comparing the centroid of every object in the binary mask to the manual tracking coordinates and removing every object but the object closest to the tracking coordinate. Finally, a watershed filter was applied to remove cells which transiently collided with the selected cell. If the selected cell could not be satisfactorily separated from neighbors it was discarded in the dataset. All shape and motion descriptors were created using the properties extract from this centroid using Matlab’s regionprops function.

These binary masks were then used to create kymographs of membrane intensity over time. The boundary of the binary mask was identified and then smoothed using a moving average filter. Then, the membrane intensity (**I_M_**) at equally spaced points along the membrane was determined by taking a 3 µm long, 1-pixel wide line scan perpendicular to the boundary and finding the maximum intensity along that boundary. I_M_ was equal to the sum of that brightest pixel along and any pixel 250 nm in either direction of the pixel along the line scan. The number of pixels (**n_pix_**) summed was saved for later. Then, cytosolic intensity (**I_C_**) was calculated by performing a binary erosion to original image mask to remove the membrane and then taking the average intensity inside that eroded mask. A background intensity (**I_bg_**) was acquired by manually selecting a 1 µm box outside of a cell and taking the average intensity in that box. Finally, the background-corrected membrane-to-cytosol ratio (**M/C**) at each point on the membrane was acquired using this formula: 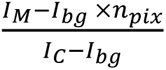. This value was then used for kymograph display.

To calculate the average M/C ratio at each timepoint, each line in a kymograph, excluding black space, was averaged. Alternatively, for dim biosensors with variable expression levels, (**Fig. 1C, 1I**), the percentage of pixels in each line above an intensity threshold was calculated. This threshold was determined by hand-identifying 3 separate patches in the image and finding the value 10 percentile points lower than the average of those three pixels in the distribution of all kymograph intensities. To average over an entire movie (**Fig. 1C**), the intensity at each timepoint recorded was simply averaged. To create a plot of average intensity or area over time, all traces were synchronized to the specified perturbation and then all values were placed in time “bins” relative to 0 where 0 is the rightmost edge of a bin and then averaged. Bin width was determined by the rate of acquisition but was generally 30 seconds. In all cases, intensities were then normalized to the average intensity from -1 minute to 0.

The calculate the membrane intensity in TIRF images (**Fig. S1B, S2B, 2D-E, 5E-F**), a mask of the entire cell was first created at every timepoint using Otsu’s method on a background-subtracted image similar to confocal images. For the average intensity (**Fig. 2D-E, S2B**), all pixels in the original image within this mask were averaged and then background-corrected by subtracting the average intensity of a 1 µm box outside the cell. For data with multiple timepoints (**Fig. 2D-E**), intensities were then normalized to the average intensity from -1 minute to 0 prior to CK666 addition. To obtain the percentage of area occupied by a wave (**Fig. S1B, 5E-F**), a wave mask was created using a threshold calculated from only intensities inside the cell mask on a frame in the movie where there is a clear active wave. The percentage was calculated by dividing the area of the wave mask by the area of the cell mask and multiplying by 100. All cells were selected based on criteria of cell health, biosensor expression, SSPB recruitment (where applicable) and wave formation prior to perturbation.

To calculate the average biosensor patch number (**Fig. 3B, 3E**), the number of patches was identified manually at every timepoint. The total number of patches over an interval (for example, all timepoints prior to CK666 addition) was then divided by the total time of that interval.

To calculate the angle of protrusion formation relative to SSPB recruitment (Fig. 5H), in FIJI, a line was drawn from the center of the cell to the center of the stimulation region and then another from the center of the cell to center of the first protrusion formed after stimulation. The angle between these two lines was calculated using each of their angles relative to the x axis of the image.

To calculate the RBD extinction from light (**Fig. 5K**), a linear kymograph was created by drawing a line through the region of light stimulation and through the wave targeted by the stimulation in FIJI and then using the reslice tool. The extinction rate is calculated by drawing one line along the border of the RBD region 30 s (10 timepoints) before stimulation and another along the border 30 s after. The height of this line (µm traveled) divided by the width (time) indicates the rate the RBD patch is displacing from the region of stimulation.

To calculate the cytoplasmic intensity of RBD in cAMP MHCKC expressing cells with or without rapamycin (**Fig. S2D**), the cytoplasm of a cell was identified manually using the freehand selection tool in FIJI. The intensity in this selection was averaged and then background-corrected by subtracting the average intensity of a 1 µm box outside the cell. Intensities were then normalized to the average intensity at 0.

### Simulations

The simulations are based on a model, previously described, in which three interacting species, RasGTP, PIP2, and PKB, form an excitable network ^24^. The concentrations of each of these molecules is described by stochastic, reaction-diffusion partial differential equations:

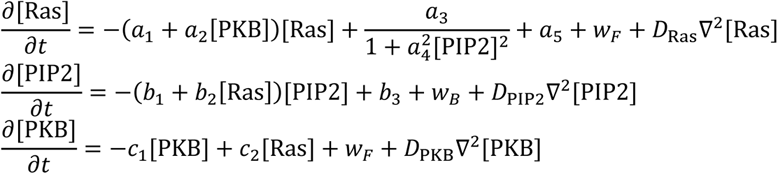

In each of these equations, the final term represents the diffusion of the species, where *D*_∗_ is the respective diffusion coefficient and ∇^2^ is the spatial Laplacian (in one or two dimensions). The second-to-last terms represent the molecular noise. Our model assumes a Langevin approximation in which the size of the noise is based on the reaction terms^85^. For example, in the case of PKB, the noise is given by

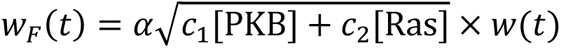

where *w*(*t*) is a zero mean, unit variance Gaussian, white noise process. In the simulations, the size of this noise was adjusted with the empirical parameter α.

In addition to the excitable dynamics described above, we incorporated two other terms. The first represents feedback from branched actin onto Ras. Since the cytoskeleton is not directly modeled, we model the origin of this feedback from PKB, which is upstream to the cytoskeleton^24^.

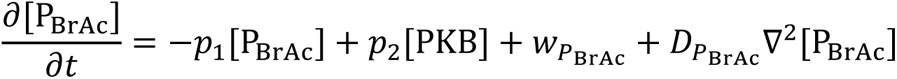

The second feedback loop accounts for the inhibitory regulation by actomyosin on Ras that originates from back molecules, such as PIP2.

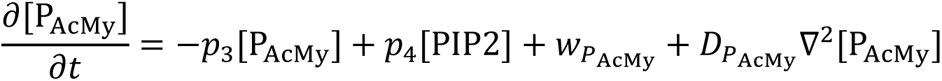

Together, they modify the equation for Ras:

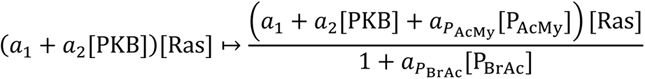

having the effect of increasing front and back contributions. In both equations, we include noise and diffusion terms.

Simulations were run on MATLAB 2023a (Mathworks) on custom-code based on the Ito solution in the Stochastic Differential Equation toolbox (http://sdetoolbox.sourceforge.net). Two-dimensional simulations were used to recreate the observed wave patterns of larger electrofused cells, and so assume a grid 40 µm × 40 µm with a spacing of 0.4 µm × 0.4 µm per grid point (*i.e*., 100 × 100 points) and zero flux boundary conditions. The one-dimensional simulations aim to recreate the membrane fluorescence observed in single-cell confocal images. The dimension is, therefore, smaller, assuming a cell radius of 5 µm and a spacing of 0.25 µm, resulting in 2μ × 5/0.25 ≈ 126 points along the perimeter, and periodic boundary conditions.

**Table 1:**
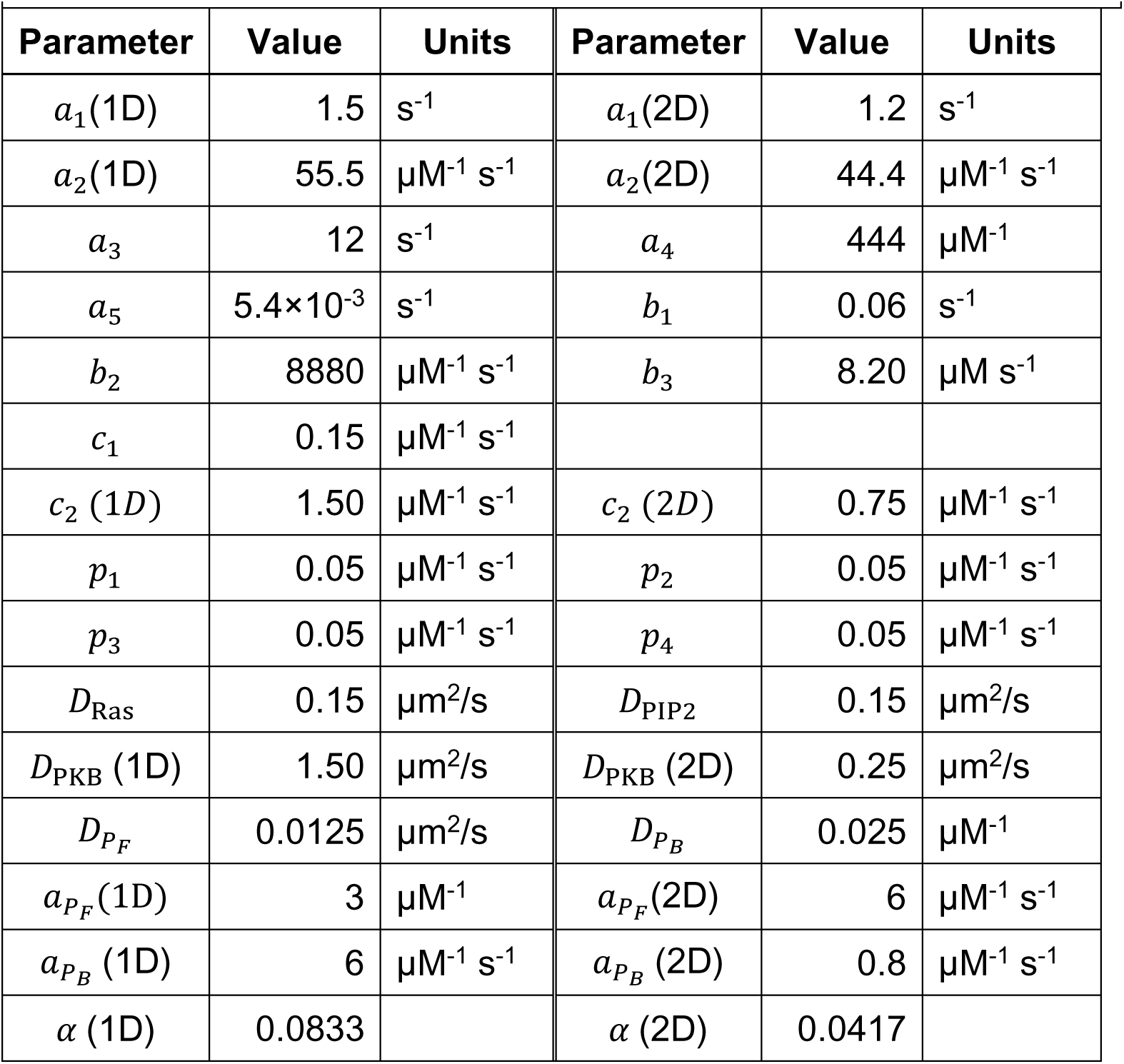
Parameters used in simulation.

| Table 1: Parameters used in simulation |  |  |  |  |  |
| --- | --- | --- | --- | --- | --- |
| Parameter | Value | Units | Parameter | Value | Units |
| $a_1(1D)$ | 1.5 | $s^{-1}$ | $a_1(2D)$ | 1.2 | $s^{-1}$ |
| $a_2(1D)$ | 55.5 | $\mu M^{-1} s^{-1}$ | $a_2(2D)$ | 44.4 | $\mu M^{-1} s^{-1}$ |
| $a_3$ | 12 | $s^{-1}$ | $a_4$ | 444 | $\mu M^{-1}$ |
| $a_5$ | $5.4 \times 10^{-3}$ | $s^{-1}$ | $b_1$ | 0.06 | $s^{-1}$ |
| $b_2$ | 8880 | $\mu M^{-1} s^{-1}$ | $b_3$ | 8.20 | $\mu M s^{-1}$ |
| $c_1$ | 0.15 | $\mu M^{-1} s^{-1}$ | | | |
| $c_2(1D)$ | 1.50 | $\mu M^{-1} s^{-1}$ | $c_2(2D)$ | 0.75 | $\mu M^{-1} s^{-1}$ |
| $p_1$ | 0.05 | $\mu M^{-1} s^{-1}$ | $p_2$ | 0.05 | $\mu M^{-1} s^{-1}$ |
| $p_3$ | 0.05 | $\mu M^{-1} s^{-1}$ | $p_4$ | 0.05 | $\mu M^{-1} s^{-1}$ |
| $D_{Ras}$ | 0.15 | $\mu m^2/s$ | $D_{PIP2}$ | 0.15 | $\mu m^2/s$ |
| $D_{PKB}(1D)$ | 1.50 | $\mu m^2/s$ | $D_{PKB}(2D)$ | 0.25 | $\mu m^2/s$ |
| $D_{PF}$ | 0.0125 | $\mu m^2/s$ | $D_{PB}$ | 0.025 | $\mu M^{-1}$ |
| $a_{PF}(1D)$ | 3 | $\mu M^{-1}$ | $a_{PF}(2D)$ | 6 | $\mu M^{-1} s^{-1}$ |
| $a_{PB}(1D)$ | 6 | $\mu M^{-1} s^{-1}$ | $a_{PB}(2D)$ | 0.8 | $\mu M^{-1} s^{-1}$ |
| $\alpha(1D)$ | 0.0833 | | $\alpha(2D)$ | 0.0417 | |

To determine the effect of these firings on cell movement, we follow an approach that simulates the movement of cells using a center-of-mass approximation. Using the spatially dependent level of activity, shown in the kymographs in **Fig. 6B**, we generated a series of force vectors normal to the cell surface. The vector sum of all these vectors was used to obtain a net protrusive force. After scaling this force so that it is in the range of experimentally observed protrusive pressures (0.5–5 nN/m^2^), we used it to push a viscoelastic model of *Dictyostelium* mechanics^86^. In this model, the net stress in the *x*-direction: σ (the direction of the gradient) alters the center-of-mass position (CM) through the following dynamics:

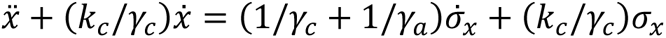

with a similar equation for the displacement in the *y*-direction (CM). Parameter values adopted from previous work^87^ are provided in the following table:

**Table 2:**
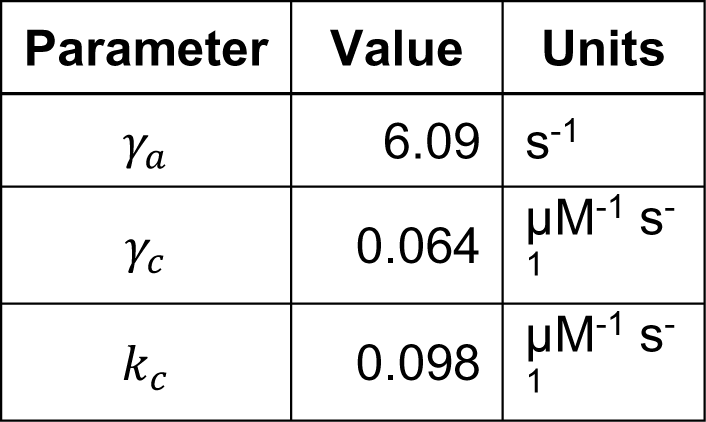
Parameters from Yang et al.

## Contributions

JK, PND, and DNR conceived the experiments. JAK performed the experiments and most analysis of experimental data. AH and PAI assisted with image analysis. PB and PA. developed the computational model and performed simulations. The manuscript was primarily written by JK and PND. with some initial writing generated by ChatGPT3 (OpenAI). Text pertaining to computational modeling was written by PB and PAI. DNR assisted in editing the paper.

## Acknowledgements

We thank members of the Peter Devreotes, Doug Robinson, Pablo Iglesias, and Miho Iijima labs for their helpful feedback and discussion. We thank the Iijima and Chuan-Hsiang Huang lab for supplying useful constructs. This work was supported by NIH grant R35 GM118177 (PND), DARPA HR0011-16-C-0139 (PAI/PND/DNR), AFOSR MURI FA95501610052 (PND), and NIH grant S10 OD016374 (S. Kuo, JHU Microscope Facility).

