## Supplemental Figures and Legends for "Complementary Cytoskeletal Feedback Loops Control Signal Transduction Excitability and Cell Polarity"

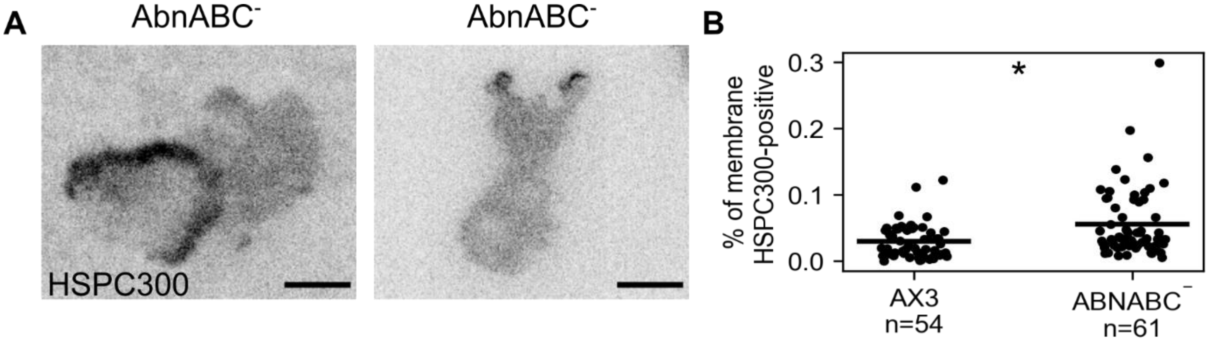


**Figure S1: WAVE activity is increased in Actobindin mutants.**

**(A)** TIRF imaging of WAVE complex unit HSPC300 (EGFP-HSPC300) in wild-type (AX3) or *actobindinABC* triple knockout (*abnABC*⁻) *Dictyostelium* cells. Images represent a single timepoint in a longer movie. *Scale bars = 5 µm.* **(B)** Individual (dots) and average (lines) percentages of the membrane area with HSPC300 localization significantly above background in AX3 and *abnABC*⁻ cells*. n = cells, *= p< 0.05*.

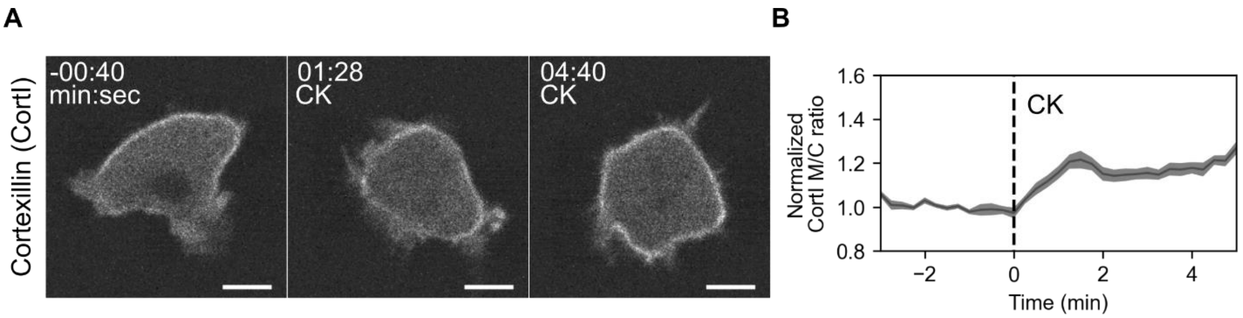
A
B
Cortexillin (Cortl)
-00:40
min:sec
01:28
CK
04:40
CK
Normalized
Cortl M/C ratio
1.6
1.4
1.2
1.0
0.8
CK
−2
0
2
4
Time (min)

8

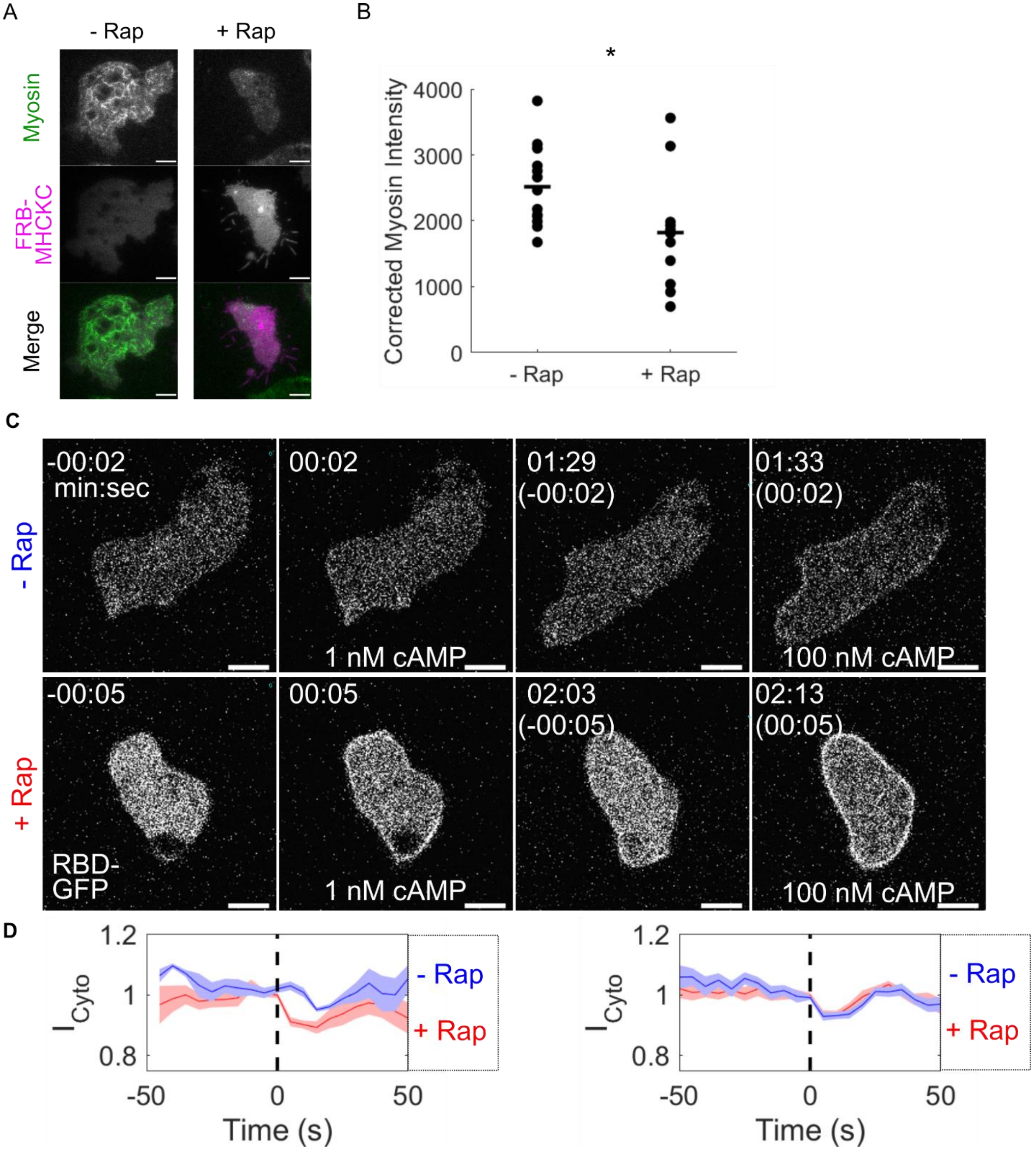
A
- Rap
+ Rap
Myosin
FRB-
MHCKC
Merge
B
*
Corrected Myosin Intensity
4000
3000
2000
1000
0
- Rap
+ Rap
C
- Rap
+ Rap
-00:02
min:sec
00:02
1 nM cAMP
01:29
(-00:02)
01:33
(00:02)
100 nM cAMP
-00:05
RBD-
GFP
00:05
1 nM cAMP
02:03
(-00:05)
02:13
(00:05)
100 nM cAMP
D
1.2
1
0.8
$I_{Cyto}$
-50
0
50
Time (s)
- Rap
+ Rap

9

**Figure S2. MHCKC disassembles myosin filaments and lowers the threshold for chemotactic stimulation**

**(A)** TIRF imaging comparing GFP-Myosin II assembly (Myosin II) and MHCKC (mCherry-FRB-MHCKC) membrane recruitment in *myosin II* null (myoII$^-$) *Dictyostelium* cells with (+ Rap) or without (- Rap) rapamycin. Cells are also expressing an unlabeled membrane-localized FKBP domain (cAR1-2xFKBP). **(B)** Individual (dots) and average (lines) of Myosin intensity on the basal membrane of Myo$^-$ cells with (+ Rap) or without (- Rap) MHCKC recruitment. *n = 13 cells (- rap) or 12 cells (+ rap)*. **(C)** Scanning confocal imaging of Ras activation (RBD-GFP) in developed *Dictyostelium* cells with (+ Rap) or without (- Rap) MHCKC recruitment. MHCKC recruitment in + Rap was confirmed but not shown here. Cells were subjected to a low dose of cAMP (1 nM) at t=00:00 and a high dose (100 nM) at t = (00:00). **(D)** Average (lines) and SEM (shaded areas) of cytoplasmic RBD intensity ($I_{Cyto}$) in cells with (+ Rap) or without (- Rap) MHCKC recruitment after a low dose of cAMP (1 nM, left) or a high dose (100 nM, right). A drop in RBD intensity in the cytoplasm is caused by translocation to the membrane and therefore an increase in Ras activity. cAMP addition occurs at t=0 (dashed line). *n = 19 cells (+ rap) and 21 cells (-rap) Scale bars = 5 µm; time is in min:sec*.

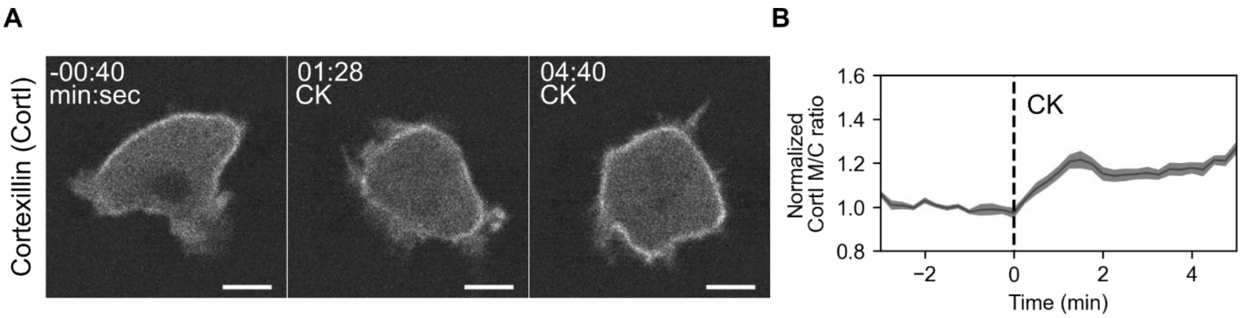


**Figure S3. Arp 2/3 inhibition increases the abundance of cortical actin.**

**(A)** Spinning disk confocal imaging of actin-actin, actin-membrane, and actin-myosin crosslinker GFP-Cortexillin I (CortI) in wild type (AX3) *Dictyostelium* cells before and after addition of Arp2/3 inhibitor CK666. *t = 00:00 indicates CK666 addition (CK); scale bars = 5 µm.* **(B)** Average (line) and SEM (shaded area) of the normalized Cortexillin I membrane-to-cytoplasm intensity ratio in cells before and after CK666 addition (dashed line, t = 0). *n = 27 cells*.

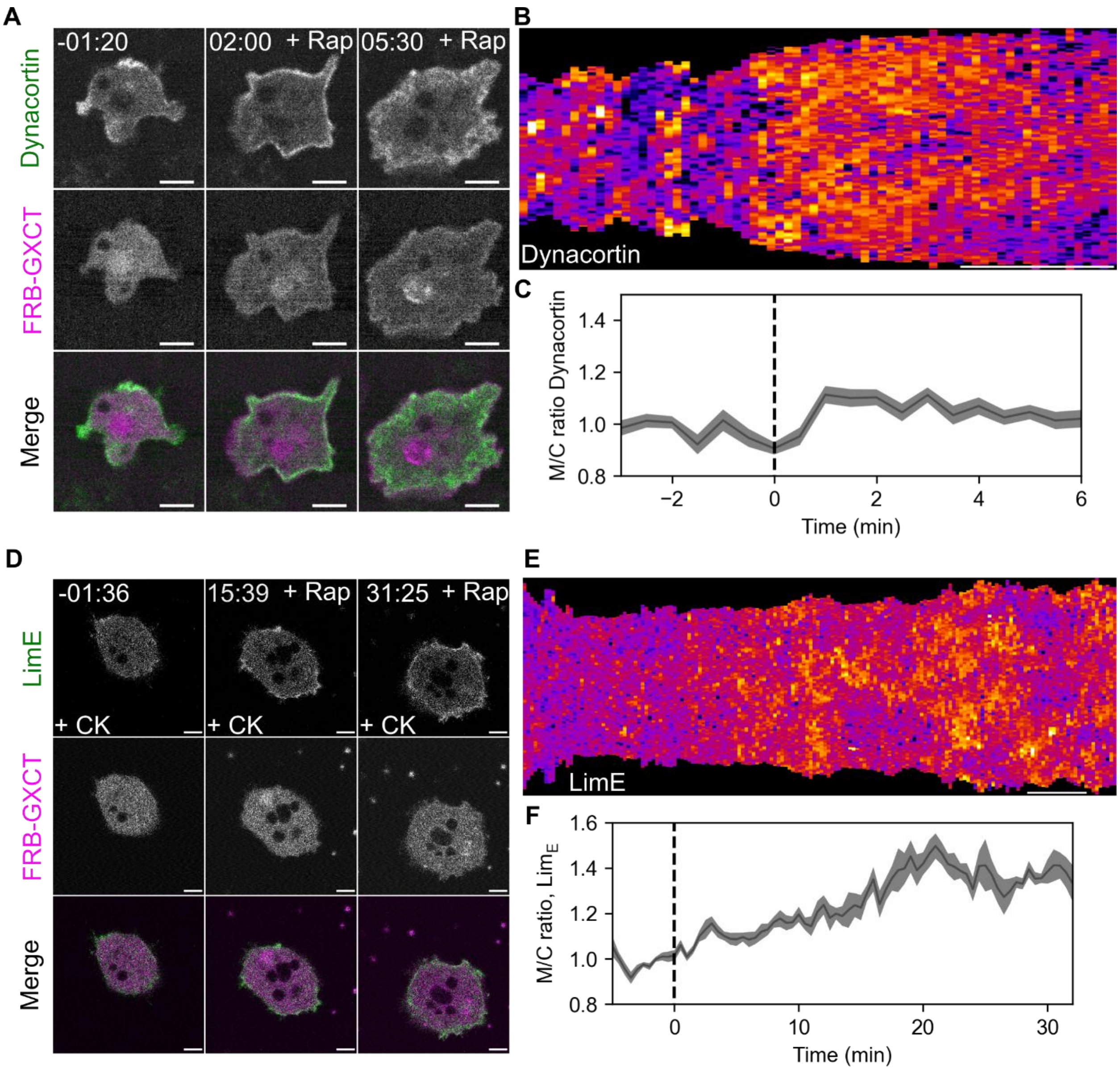


**Figure S4: RacE activation triggers Arp2/3-independent actin polymerization and recruits actin crosslinkers.**

**(A)** Scanning Confocal imaging of GXCT recruitment and actin crosslinker dynacortin (EGFP-Dynacortin) in wild type (AX3) *Dictyostelium* cells. *t = 00:00 indicates rapamycin addition.* **(B)** Membrane kymograph of dynacortin from the movie in (A). *Scale bar = 3 minutes.* **(C)** Average (line) and SEM (shaded area) membrane-to-cytosol ratio of dynacortin before and after GXCT recruitment by rapamycin addition (dashed line, t = 0). *n = 7 cells.* **(D)** Scanning Confocal imaging of GXCT recruitment and polymerizing actin (LimE$_{\Delta Coil}$-EGFP) in wild type (AX3) *Dictyostelium* cells treated with the Arp2/3 inhibitor CK666. *t = 00:00 indicates rapamycin addition.* **(E)** Membrane kymograph of LimE from the movie in (D). *Scale bar = 5 minutes.* **(F)** Average (line) and SEM (shaded area) membrane-to-cytosol ratio of LimE before and after GXCT recruitment by rapamycin addition (dashed line, t = 0) in cells treated with CK666. *n = 7 cells.*

**Movie S1:** Timelapse scanning confocal video of Ras activation (RBD-GFP) in wild type (AX3, left) and *actobindin* triple-null (*abnABC*$^{-}$, right) *Dictyostelium* cells. Images acquired every 7 (left) or 6 (right) seconds. Frame rate is 12 frames per second. *Scale bar = 5 µm; time is in min:sec. Related to figure 1.*

**Movie S2:** Timelapse scanning confocal video of Myosin Heavy Chain Kinase C (mCherry-FRB-MHCKC) recruitment in wild type (AX3) *Dictyostelium* cells. Cells are also expressing an unlabeled membrane-localized FKBP domain (cAR1-2xFKBP). Images acquired every 30 seconds. Frame rate is 20 frames per second. *Scale bar = 5 µm; time is in min:sec. Time t = 00:00 indicates rapamycin addition. Related to figure 1.*

**Movie S3:** Timelapse scanning confocal video of Myosin Heavy Chain Kinase C (mCherry-FRB-MHCKC, magenta) recruitment and Ras activation (RBD-EGFP) in wild type (AX3) *Dictyostelium* cells. Cells are also expressing an unlabeled membrane-localized FKBP domain (cAR1-2xFKBP). Images acquired every 15 seconds. Frame rate is 20 frames per second. *Scale bar = 5 µm; time is in min:sec. Time t = 00:00 indicates rapamycin addition. Related to figure 1.*

**Movie S4:** Timelapse scanning confocal video of polymerizing actin ($LimE_{\Delta Coil}$-mCherry) and $PIP_3$ levels ($PH_{CRAC}$-YFP) in wild type (AX3) *Dictyostelium* cells before and after treatment with the Arp 2/3 inhibitor CK666 (CK). Images acquired every 15 seconds. Frame rate is 15 frames per second. *Scale bar = 5 µm; time is in min:sec. Time t = 00:00 indicates CK666 addition. Related to figure 2.*

**Movie S5:** Timelapse TIRF video of activated Ras (RBD-EGFP) in electrofused ("giant") wild type *Dictyostelium* cells before and after CK666 addition and subsequently latrunculin addition. Cells are incubated in caffeine to raise basal activity levels. Images acquired every 26 seconds. Frame rate is 12 frames per second. *Scale bar = 5 µm; time is in min:sec. Time t = 00:00 indicates CK666 addition. Related to figure 2.*

**Movie S6:** Timelapse scanning confocal video of PIP3 levels ($PH_{AKT}$) in human neutrophil-like (dHL60) cells before and after the addition of CK666 addition and subsequently latrunculin addition. Images acquired every 12 seconds. Frame rate is 15 frames per second. *Scale bar = 5 µm; time is in min:sec. Time t = 00:00 indicates CK666 addition. Related to figure 2.*

**Movie S7:** Timelapse scanning confocal video of PIP3 levels ($PH_{CRAC}$-YFP) in *myosin II*-null (*myoII*$^{-}$) *Dictyostelium* cells before and after CK666 addition. Images acquired every 9 seconds. Frame rate is 12 frames per second. *Scale bar = 5 µm; time is in min:sec. Time t = 00:00 indicates CK666 addition. Related to figure 3.*

**Movie S8:** Timelapse scanning confocal video of Myosin Heavy Chain Kinase C (mCherry-FRB-MHCKC, magenta) recruitment and Ras activation (RBD-EGFP) in wild type (AX3) *Dictyostelium* cells before and after CK666 addition and subsequently rapamycin addition. Cells are also expressing an unlabeled membrane-localized FKBP domain (cAR1-2xFKBP). Images acquired every 15 seconds. Frame rate is 16 frames per second. *Scale bar = 5 µm; time is in min:sec. Time t = 00:00 indicates CK666 addition. Related to figure 3.*

**Movie S9:** Timelapse scanning confocal video of RacE-GEF (mCherry-FRB-GXCT$_{\Delta NT}$) membrane recruitment in wild type (AX3) *Dictyostelium* cells. Cells are also expressing an unlabeled membrane-localized FKBP domain (cAR1-2xFKBP). Images acquired every 15 seconds. Frame rate is 15 frames per second. *Scale bar = 5 µm; time is in min:sec. Time t = 00:00 indicates rapamycin addition. Related to figure 4.*

**Movie S10:** Timelapse scanning confocal video of RacE-GEF (mCherry-FRB-GXCT$_{\Delta NT}$) membrane recruitment and polymerizing actin (LimE$_{\Delta Coil}$-EGFP) in wild type (AX3) *Dictyostelium* cells. Cells are also expressing an unlabeled membrane-localized FKBP domain (cAR1-2xFKBP). Images acquired every 10 seconds. Frame rate is 15 frames per second. *Scale bar = 5 µm; time is in min:sec. Time t = 00:00 indicates rapamycin addition. Related to figure 4.*

**Movie S11:** Timelapse scanning confocal video of RacE-GEF (mCherry-FRB-GXCT$_{\Delta NT}$) membrane recruitment and Ras activation (RBD-EGFP) in wild type (AX3) *Dictyostelium* cells. Cells are also expressing an unlabeled membrane-localized FKBP domain (cAR1-2xFKBP). Images acquired every 10 seconds. Frame rate is 15 frames per second. *Scale bar = 5 µm; time is in min:sec. Time t = 00:00 indicates rapamycin addition. Related to figure 4.*

**Movie S12:** Timelapse scanning confocal video of RacE-GEF (tagRFP-SSPB-GXCT$_{\Delta NT}$) optical membrane recruitment in wild type (AX3) *Dictyostelium* cells. Cells are also expressing an unlabeled membrane-localized iLID domain (N150-iLID). Images acquired every 12 seconds. Frame rate is 5 frames per second. *Scale bar = 5 µm; time is in min:sec. Time t = 00:00 indicates blue light exposure. Related to figure 5.*

**Movie S13:** Timelapse TIRF video of activated Ras (RBD-emiRFP670) in electrofused ("giant") wild type *Dictyostelium* cells before, during, and after tagRFP-SSPB-GXCT$_{\Delta NT}$ recruitment. The cell begins with no light stimulation, is exposed to uniform 488 nm light (+ 488) which is then removed as indicated. Cells are also expressing an unlabeled membrane-localized iLID domain (N150-iLID). Images acquired every 9 seconds. Frame rate is 10 frames per second. *Scale bar = 5 µm; time is in min:sec. Time t = 00:00 indicates blue light exposure. Related to figure 5.*

**Movie S14:** Timelapse scanning confocal video of RacE-GEF (SSPB-GXCT$_{\Delta NT}$) or negative control (SSPB) local membrane recruitment in wild type (AX3) *Dictyostelium* cells with or without pre-incubation with Arp2/3 inhibitor CK666. Blue box indicates the region of blue light exposure. Cells are also expressing an unlabeled membrane-localized iLID domain (N150-iLID). Images acquired every 6 seconds. Frame rate is 6 frames per second. *Scale bar = 5 µm; time is in min:sec. Time t = 00:00 indicates blue light exposure. Related to figure 5.*

**Movie S15:** Timelapse TIRF video of activated Ras (RBD-emiRFP670) in electrofused ("giant") wild type *Dictyostelium* cells before and after RacE-GEF (SSPB-GXCT$_{\Delta NT}$) local membrane recruitment. Cells are also expressing an unlabeled membrane-localized iLID domain (N150-iLID). Blue box indicates the region of blue light exposure. Movie on the right is a 3X scaled zoom of the area of interest on the left. Images acquired every 3 seconds. Frame rate is 10 frames per second. *Scale bars = 5 μm; time is in min:sec. Time t = 00:00 indicates blue light exposure. Related to figure 5.*

**Movie S16:** This timelapse 2D simulation depicts Ras/PIP2 waves in Dictyostelium cells, emphasizing the effects of CK666 and Latrunculin treatment on wild-type cells. Initial 249 seconds show wild-type behavior with both feedback loops active. From 250 to 449 seconds, branched actin feedback is selectively deactivated to model CK666's effects on actin dynamics. In the last 200 seconds, the actomyosin feedback loop is disabled, simulating Latrunculin's additional impact on CK666-treated cells. The simulation accurately represents spatial dimensions, with a length of 40 microns along each axis, mirroring typical Dictyostelium cell dimensions.
